# We shall meet again - Genomics of historical admixture in the sea

**DOI:** 10.1101/2020.05.01.069740

**Authors:** Xueyun Feng, Juha Merilä, Ari Löytynoja

**Affiliations:** Ecological Genetics Research Unit, Organismal and Evolutionary Biology Research Programme, Faculty of Biological and Environmental Sciences, 00014 University of Helsinki, Finland; Institute of Biotechnology, University of Helsinki, 00014 Helsinki, Finland

**Keywords:** adaptation, admixture, gene flow, effective population size, recombination rate, secondary contact

## Abstract

We studied the impact of genetic introgression in evolution and on evolutionary studies with whole-genome data from two divergent lineages of sticklebacks. Our results reveal that the hybrid zone between the lineages ranges across the entire Baltic Sea and parts of the North Sea with the foreign ancestry decreasing with increasing distance to the source population. Introgression has also penetrated currently isolated freshwater populations. We identified footprints of selection on regions enriched for introgressed variants, suggesting that some of the introgression has been adaptive. However, the levels of introgression were in general negatively correlated with the recombination rate, suggesting that the introgression has been largely neutral and adaptive ancestral standing variation likely had a more important role in shaping the genomic landscape. Our results further suggest that overlooked introgression can mislead analyses of local adaptation and phylogenetic affinities, highlighting the importance of accounting for introgression in studies of local adaptation.

## Introduction

Introgression is an important evolutionary process that transfers genetic variation between divergent lineages. Whole-genome scale analyses have shown introgression to be an important and pervasive evolutionary force shaping the genomes of many organisms (e.g. Lamichhaney et al. 2018; Oziolor et al. 2019; Suarez-Gonzalez et al. 2018), including humans (Huerta-Sanchez et al. 2014; Sankararaman et al. 2014, 2016; Racimo *et al.* 2015). Although generally a homogenizing process, there is also accumulating evidence of introgression fuelling speciation and local adaptation (Lamichhaney et al. 2018; Marques, Meier, and Seehausen 2019; Suarez-Gonzalez et al. 2018; Hedrick 2013; Oziolor et al. 2019). Because of limitations in both analytical methods and sampling, ancient introgression is often neglected in population genetic studies, and its importance, consequences, and contributions in adaptive evolution and speciation of non-model organisms are poorly understood.

Genomic studies of contemporary hybrids have shown that the rate of introgression is highly variable among species (Martin *et al.* 2015, 2019; Malinsky *et al.* 2018) and populations (Skoglund *et al.* 2015; Kuhlwilm *et al.* 2016), and the extent of introgressed ancestry is unevenly distributed across the genome (Sankararaman *et al.* 2014, 2016; Martin *et al.* 2019; Edelman *et al.* 2019; Stankowski *et al.* 2019). The underlying mechanisms shaping this heterogeneous distribution are not well understood. Generally, introgressed alleles are regarded to have a negative fitness effect when introduced into new genomic backgrounds (Martin and Jiggins 2017; Bay *et al.* 2019) and, according to the Dobzhansky-Muller model of hybrid incompatibility (Masly *et al.* 2007, Bomblies *et al.* 2007, Lee *et al.* 2008), long-term negative selection on incompatible loci may create ‘deserts’ of introgression in the genome (Sankararaman *et al.* 2014, 2016). Genetic architecture and constraints also play a role, and genomic regions characterized by higher gene density and(or) low recombination rate are expected to show lower levels of introgression (Barton and Bengtsson 1986). On the other hand, the populations involved in introgression can differ in their dynamics of deleterious variation and genetic load, affecting the patterns of selection and subsequent distribution of introgressed ancestry (Henn *et al.* 2016, Peischl and Excoffier 2015). Low effective population size (*N_e_*) and strong drift may allow weakly deleterious alleles to accumulate in one population and thus reduce the relative fitness of the genetic material introgressed to the other population. The high genetic load has been proposed to be the main driver of selection against Neanderthal introgression in non-African humans (Harris & Nielsen 2016; Juric, Aeschbacher and Coop 2016). Large differences in historical *N_e_* among hybridizing populations have likely been common, but it remains largely unknown how such factors shape the genomic landscape of introgression.

Although introgressed alleles are generally thought to have negative effects on fitness and be selected against, some introgressed alleles can be adaptive and rapidly spread in the recipient population (Hedrick 2013; Suarez-Gonzalez *et al.* 2018). Such adaptive introgression is expected to leave a unique footprint of enriched introgressed ancestry. For instance, altitude adaptation in Tibetan humans (Huerta-Sánchez *et al.* 2014), the beak shape of Darwin’s finches (Lamichhaney *et al.* 2018), and mimicry patterns of a butterfly (Enciso-Romero *et al.* 2017) are examples of adaptive traits introgressed from a related population or species. Several recent studies have taken a genome-wide approach to find correlated signatures of selection favouring introgressed genetic variations (e.g. Racimo *et al* 2017; Richards and Martin, 2017; Oziolor *et al.* 2019; Enciso-Romero *et al.* 2017). Most of these studies have focused on terrestrial organisms, and little is known about the possible role of adaptive introgression in the marine environment (but see: Richards and Martin 2017; Oziolor *et al.* 2019), where physical barriers to gene flow are minimal, and the effects of genetic drift are reduced due to large population sizes (Palumbi 1994; Jokinen *et al.* 2019). The aim of this study was to investigate the extent and mechanisms of introgression between two ancient lineages of nine-spined sticklebacks (*Pungitius pungitius*) following their secondary contact in northern Europe. To this end, we obtained Whole-Genome Resequencing (WGR) data from 289 individuals from 14 populations, along with an outgroup species, and estimated the levels of introgression across different populations and genomic regions to characterize the hybrid zone. Our primary goals were to 1) confirm the introgression and determine its geographic distribution, 2) understand the evolutionary forces that have shaped the genomic landscape of introgression, 3) identify the genomic regions where introgression likely has been facilitated by positive natural selection favouring introgressed genetic elements, and 4) investigate the effects of genetic introgression on typical phylogenomic and population genetic analyses. Our results show that ancient introgression has penetrated deep into the distribution range of these previously separated lineages, and contributed to local adaptation. However, we find that the proportion of foreign ancestry is negatively correlated with the recombination rate and suggest that this may be caused by positive selection on ancestral standing variation, with introgressed variation being largely neutral. Our findings also highlight the importances of taking historical admixture into account in population genetic analyses, including studies on the genetic basis of local adaptation.

## Results

### Whole-genome resequencing and variant calling

We resequenced 290 individuals from 14 populations (Fig. 1) to an average depth of 12.5X (range 8–36.7X). After alignment, variant calling and quality control we obtained 7,297,498 single nucleotide polymorphisms (SNPs) over the 20 autosomal linkage groups.

**Figure 1.**
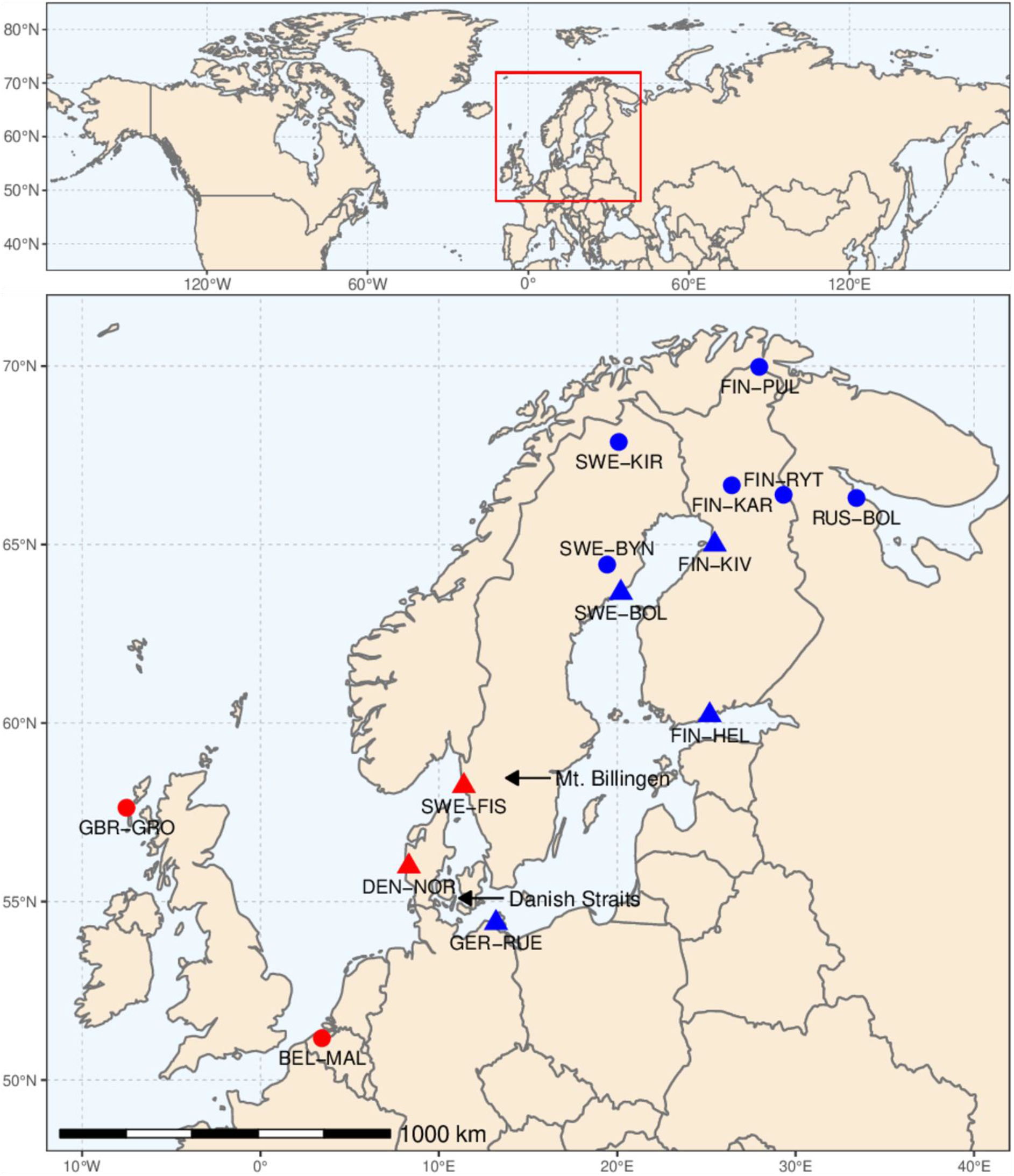
Geographic locations of sampled freshwater (●) and marine (▲) nine-spined stickleback populations. Red and blue colours refer to western (WL) and eastern (EL) lineage origin, respectively.

### Population genetic structure

We analysed the population structure using Principal Component Analysis (PCA) and ADMIXTURE analysis. In PCA, little differentiation was seen among the Eastern European Lineage (EL) populations or the northern Baltic Sea (NBS) populations (FIN-KIV, SWE-BOL, FIN-HEL), whereas the Western European Lineage (WL) populations were clearly differentiated (Fig. 2a). Principal Component (PC) 1, explaining 32.1% of the variance, differentiated the six EL populations from the WL lineage populations, and left the Baltic Sea populations intermediate between them (Fig. 2a, Supplementary Figure S1); PC2, explaining 13.5% of the variance, largely captured divergence among the WL populations. The ADMIXTURE analysis was consistent with the PC1 and showed intermediate ancestry proportions varying from 99.9% EL-like to 99.9% WL-like (Fig. 2b).

**Figure 2.**
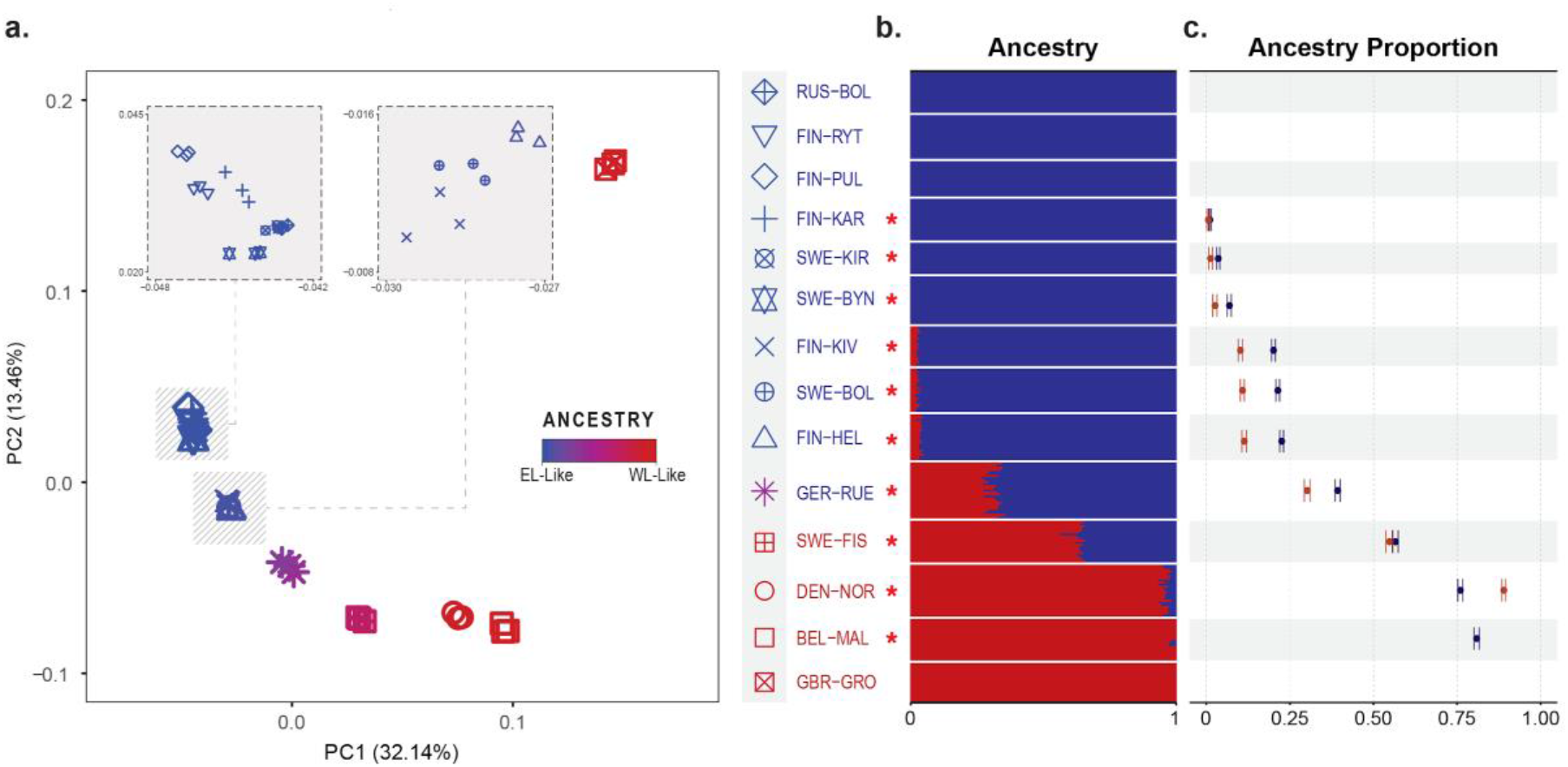
Population structure and estimated ancestry proportions of the studied populations. **a.** The first two principal components of the PCA analysis with three individuals from each nine-spined stickleback population. The colour indicates the proportion of “EL-like” ancestry as estimated with ADMIXTURE (K=2). **b.** Population genetic structure inferred with ADMIXTURE (K=2) using selected genome-wide SNPs (see Methods for details). Each row represents a population, and red (WL-like) and blue (EL-like) colours show the estimated ancestry proportions. Population labels colored in red and blue refer to the lineage of the population (WL and EL, respectively), the asterisk indicating the significance at *Z* ≥ 3 in *D*-statistic. **c.** The WL ancestry proportions estimated by *f_4_*-ratio test with 95% confidence intervals for the alpha values. Dark red and blue colours refer to direct estimation (α_WL_) or in-direct estimation (α_WL_ = 1 - α_EL_) of WL ancestry, respectively. For details about *D*-statistic and *f4-ratio* tests, see Methods and Supplementary Table S3, S4.

### Quantification of genomic introgression

The occurrence of admixture was assessed formally using *D*-statistic (Patterson *et al.* 2012) and quantified using *f_4_-ratio* test (Reich *et al.* 2009) in the *direct* manner (Petr *et al.* 2019). In line with the earlier results, the *D*-statistics indicated that the Baltic Sea populations have admixed with the WL, and the North Sea populations have admixed with the EL (Fig. 2b, Supplementary Table S3). Interestingly, we also found three pond populations (SWE-BYN, SWE-KIR, and FIN-KAR) being introgressed, although for the last two the degree of significance varied depending on the EL source population used (Fig. 2b, Supplementary Table S3). Generally, the ancestry proportions showed opposite gradients on the two sides of the Danish straits with the foreign ancestry decreasing with increasing distance from the source (Fig. 2c, Supplementary Table S4). The admixed North Sea populations (BEL-MAL, DEN-NOR, SWE-FIS) contained 19.1–43.5% and the southern Baltic Sea population (GER-RUE) 60.7% of EL ancestry. The NBS populations were similar to each other and were estimated to contain 77.4–79.9% of EL ancestry and 20.1–22.6% of WL ancestry (Fig. 2c, Supplementary Table S4). In the landlocked pond populations showing significant *D*-statistic, the levels of WL introgression were low. Using multiple EL source populations, the estimated WL ancestries varied between 0–1.1%, 0.7–3.6%, and 3.6–7.0% for FIN-KAR, SWE-KIR, and SWE-BYN, respectively (Supplementary Table S4).

We binned the genome into five categories – intergenic, coding DNA (CDS), constrained elements, introns and promoters – and examined the distribution of introgressed variation across these. Using the *direct* estimation, the three NBS populations showed significantly higher WL ancestry in promoter regions (*p* = 0.022–0.048) and significantly lower WL ancestry in constrained elements (*p* = 0.005–0.046; Supplementary Table S5). For the admixed WL populations, the EL ancestry was significantly lower in the CDS (*p* < 0.0001), promoter (*p* = 0.001–0.01), and intron (*p* = 0.0136-0.0139) categories than in intergenic regions, but no difference was seen in constrained elements (*p* = 0.476–0.512).

### Dating of introgression from demographic history

Footprints in the genome may allow dating past admixture events (Hawks 2017). With the exception of Swedish inland populations, the population-specific demographic histories inferred using SMC++ indicated that all Baltic Sea area populations experienced a rapid increase in effective population size (*N_e_*) around 5,000–7,000 ya (Fig. 3). Such a pattern can be caused by introgression introducing novel genetic variation in the recipient populations, thus increasing the coalescent *N_e_*. Assuming an average generation time of two years (DeFaveri *et al.* 2014), the increase in *N_e_* coincided with the reconnection of the current Baltic Sea and the North Sea around 10,000 years ago (ya; Fig. 3; Schwarzer *et al.* 2008). The slight differences in the timing of the inferred demographic and the known geological events might owe to the inaccuracies in the mutation rate or the generation time estimate used.

**Figure 3.**
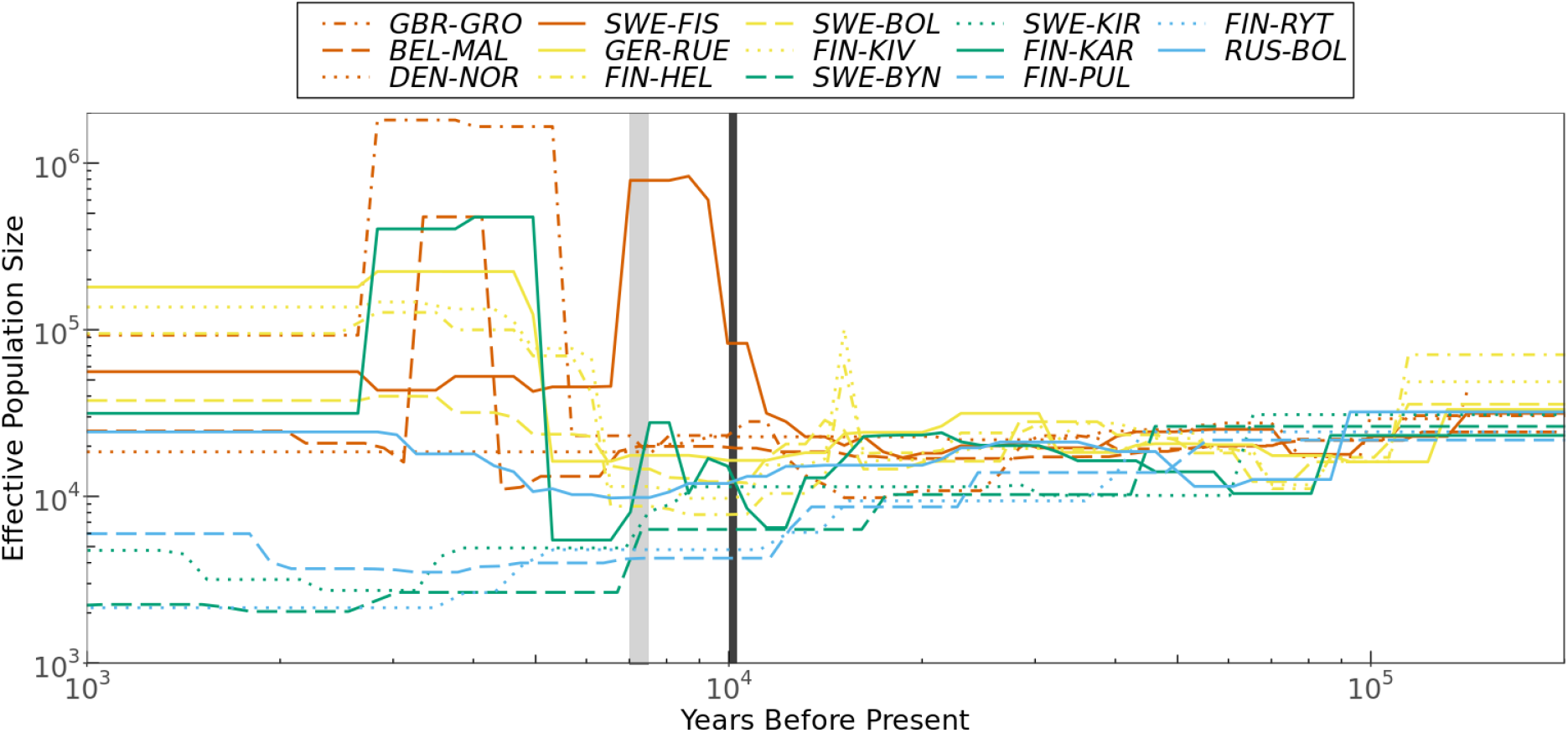
Ancestral effective population sizes (*N_e_*) of nine-spined stickleback populations as a function of time (scaled as years before present, assuming average generation time of two years). Black and grey vertical lines mark the estimated times of first (through Mt. Billingen) and second (through Danish Straits) Baltic Sea-North Sea connection, respectively. The Western Lineage, the Baltic Sea, admixed pond and the non-admixed Eastern Lineage populations are coloured by orangeish, yellowish, greenish and blueish colours, respectively.

Surprisingly, the Scottish freshwater population GBR-GRO showed the greatest increase in *N_e_*. To test whether this rapid increase was driven by introgression from EL, we used a phylogenetic approach (Methods) that classifies locus-specific phylogenetic trees according to their grouping. We found that 96.3% of the resolved trees (of total 767 trees) grouped with BEL-MAL and only 3.8% suggested EL branch-off, making the eastern introgression an unlikely explanation.

### Footprints of selection on WL introgression in Baltic Sea populations

The NBS populations seemed the most promising system to study the mechanisms and extent of genetic introgression as well as the potential adaptive nature of the introgressed variation. For greater precision, we computed the *fd* summary statistic (Martin *et al.* 2015) for 100-kb windows with 20-kb steps across the genome, replicating the analysis with three different EL references for each of the three NBS populations. Based on the FDR corrected *p*-value cut off at 0.05, and the intersection of the nine analyses, we obtained 25 putative introgression-enriched regions with lengths varying from 100–200kb (Fig. 4a, Supplementary Table S6). Shared genetic variation originating from introgression causes a decrease in *d_xy_* whereas shared variation due to ancestral polymorphism does not (Martin *et al.* 2015). Based on patterns of *d_xy_* variation as well as information on genotypes and sequencing coverage, we confirmed eight candidate regions to be introgressed segmental duplicates, five to contain introgressed deletions, and one region to be a false positive likely caused by a shared inversion (Supplementary Figure S2 and Supplementary Table S6). The last one was removed from downstream analyses.

**Figure 4.**
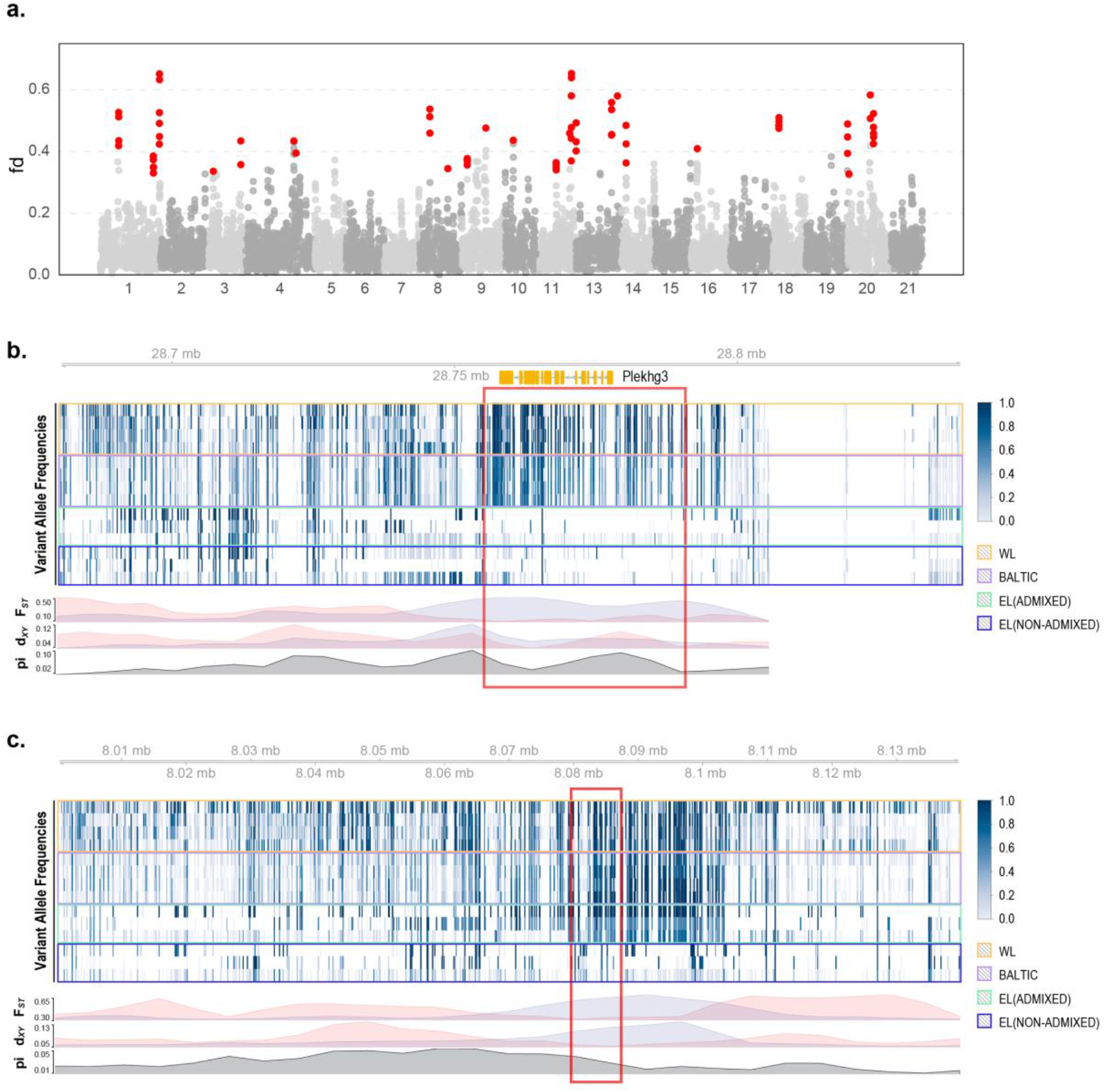
Adaptive introgression in the northern Baltic Sea populations. **a.** Manhattan plot of WL admixture as measured by *fd* across the 20 autosomal linkage groups. Each dot represents 100-kb regions and shows the mean *fd* values across nine comparisons (three northern Baltic Sea populations x three EL reference populations), red indicating regions significantly enriched for WL ancestry identified using different EL source populations (see Methods). The peak in LG18 is caused by a high-frequency inversion in EL reference populations not involved in introgression (Supplementary Figure S2). **b.** LG1:28680001-28840000 and **c.** LG11:8000001-8140000 are examples of candidate regions for adaptive introgression (AI). Shown are gene structure (coding sequences in orange), per site variant allele frequencies (heatmap), and line plots of *F_ST_*, *d_xy_* and *π* variation in northern Baltic Sea populations (10kb-windows with 5-kb step). Color frames indicate different population sets (see Supplementary Figure 2). *F_ST_* and *d_xy_* measured against RUS-BOL and GBR-GRO are shown in light blue and pink, respectively. Regions identified as adaptive introgression (see Methods) are marked with red squares.

To rule out incomplete lineage sorting (ILS), we computed the probability of observing tracts of ancestral variation as long as the 24 candidate regions (Huerta-Sánchez *et al.* 2014). Using local recombination rate estimates (1×10^-10^–1.15×10^-7^ per generation per bp) and the lower bound of divergence time (see Methods), the expected length of shared ancestral tracks ranged from 40 bp to 46,512 bp, and the probability of the observed tracks (90,059– 199,904 bp) ranged from 0 to 0.416. After excluding four tracks located in non-recombining regions (*p* > 0.05), the remaining 20 candidate regions were unlikely to be explained by ILS *(p* = 0-6.708×10^-6^), and their expected ages ranged from 97 to 29,324 generations (Supplementary Table S6).

The *U* and *Q95* tests (Racimo *et al.* 2015) distinguish adaptive introgression from neutral admixture using the number of sites uniquely shared between the donor and recipient population as well as the allele frequencies on those sites. Using these tests, we obtained seven candidate regions showing signals of selection (Supplementary Table S7) and found four of these to overlap with regions identified with the *fd* analysis. These four regions were chosen as candidates for adaptive introgression (AI). Integrating information from *F_ST_*, *d_xy_*, π, variant allele frequencies, sequencing coverage, and SNP information from *U* and *Q*95 test results, three genes were identified as targets of adaptive introgression (*viz*. *ZP4, PLEKHG3, PIK3R6;* Fig. 4b, Supplementary Figure S2 and Supplementary Tables S6 and S7).

Finally, we used shared variation to assess the relative contribution of WL genetic variation to the differentiation and adaptation of the NBS populations. In total, 4,323,090 SNPs were variable among the five populations (Methods) with 1.92% (83,057) of the SNPs being fixed between WL and EL. Of the variable sites, 64.23% (2,776,743) were polymorphic in NBS, 49.4% (2,135,474) in WL, and 33.37% (1,443,005) in EL (Fig. 5a). Correcting for the different numbers of individuals, the mean nucleotide diversity (*π*) was roughly equal in NBS and WL (0.033 vs. 0.034), and slightly lower in EL (0.027). Of the SNPs variable in NBS (2,776,743), 16.3% and 11.92% were shared with EL and WL, respectively, while the remaining were either polymorphic in all groups (22.92%) or unique to NBS (48.86%; Fig. 5b, Supplementary Table S8). In NBS, 0.68% (5,305 out of 783,691) of the EL or WL-shared SNPs were found to be under positive selection: from those 90.23% were EL-shared and 9.77% WL-shared (Fig. 5c). Given that 11.92% of the NBS SNPs were classified as WL origin, selection on those (adaptive introgression) was extremely rare (0.16%, 518 out of 331,078 SNPs).

**Figure 5.**
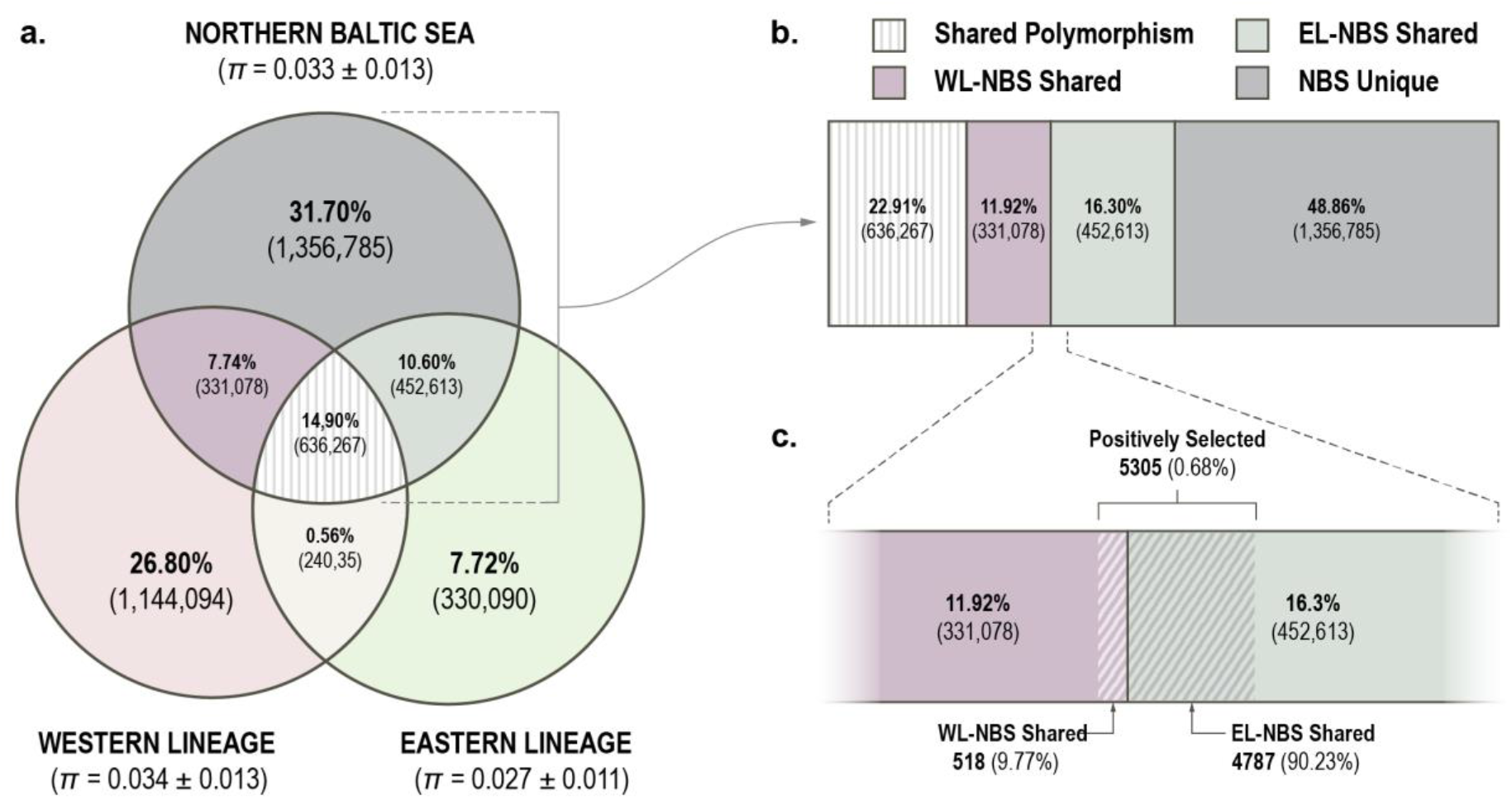
Shared polymorphisms among the northern Baltic Sea (NBS) populations and representative populations from Western Lineage (WL; GBR-GRO) and Eastern Lineage (EL; RUS-BOL). **a.** Venn diagram shows the number of polymorphic sites shared between the groups and nucleotide diversity (*π*; in parentheses, genome-wide mean and standard deviation of 100-kb window estimates) for each group. **b.** The boxes, with colours matching those in **a.**, show the relative proportion of shared and unique polymorphism in the NBS populations. **c.** The shaded area shows the proportion of WL-NBS or EL-NBS shared polymorphisms inferred to be under positive selection.

### Interaction between introgression, differentiation, and recombination rate

We observed heterogeneity in admixture proportions and differentiation across the genome (Fig. 6a). Specific patterns could be caused by the incompatibility of introgressed genetic variants interplaying with different mechanisms, for example with the local recombination rate and the removal of linked neutral variation (e.g. Schumer *et al.* 2018). On the other hand, genomic regions experiencing lower levels of recombination are expected to be more differentiated than those experiencing more recombination (Nachman and Payseur 2012). We found the population-specific correlation between the admixture proportion and the local recombination rate to be either negative or insignificant (Supplementary Table S9), though a slightly positive correlation was seen between the mean *fd* estimate and the recombination rate (Fig. 6b). Rather unexpectedly, *F_ST_* was found to be negatively correlated with recombination rate only in three of the four comparisons between ponds and the EL reference population (Table 1). Positive correlations were found in the rest of the comparisons (*r_s_* = 0.03 to 0.39, *p* < 0.05), regardless of which parental population was used in the comparison (Fig. 6c,d), or if the comparison was between the WL and EL reference populations (Fig. 6e).

**Figure 6.**
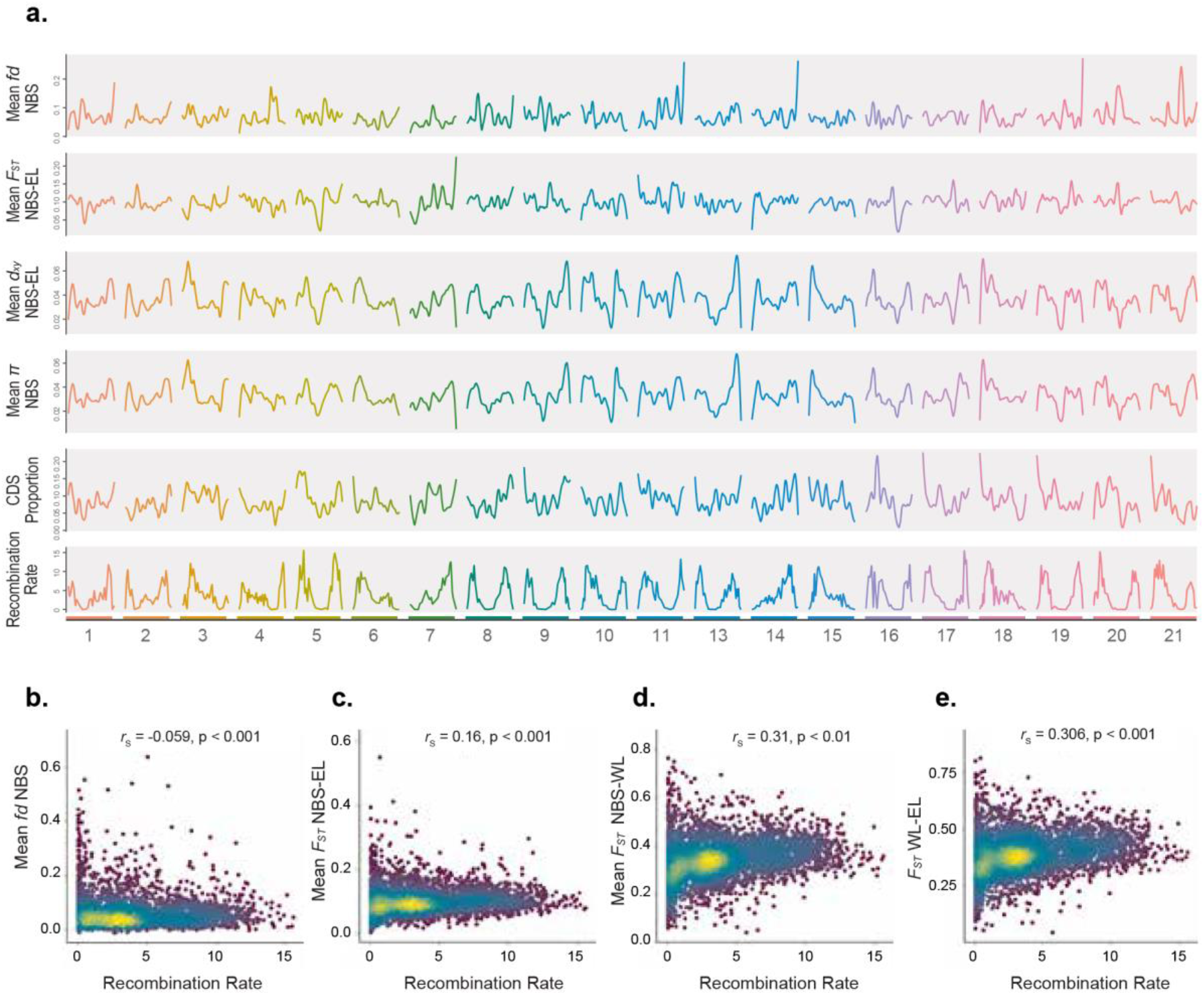
Variation in admixture proportion (*fd*), measure of population differentiation (*F_ST_*), absolute divergence (*d_xy_*), nucleotide diversity (*π*), coding sequence (CDS) density, and recombination rate across different linkage groups (LGs). **a.** Mean *fd*, *F_ST_*, *d_xy_*, π estimates of the three northern Baltic Sea (NBS) populations (FIN-HEL, SWE-BOL and FIN-KIV), CDS density, and inferred recombination rate (in cM/Mb) estimated in non-overlapping 100kb-windows. *F_ST_*, and *d_xy_* were measured against RUS-BOL. *fd*, *F_ST_*, *d_xy_*, π and CDS density estimates are smoothened (*loess*, span=0.2). Correlation between recombination rate and: **b.** mean *fd* estimates of the three NBS populations, **c.** mean *F_ST_* between the three NBS and RUS-BOL population, **d.** mean *F_ST_* between the three NBS and GBR-GRO population, and **e.** *F_ST_* between the WL (GBR-GRO) and EL (RUS-BOL) source populations. Colours indicate density of points from low (dark blue) to high (yellow).

**Table 1.** Spearman rank correlations (*r_s_*) between recombination rate and *F_ST_* estimated in 100-kb windows. Correlations are given against WL (GBR-GRO) and EL (RUS-BOL) reference populations.

|  | GBR-GRO |  | RUS-BOL |  |
| --- | --- | --- | --- | --- |
| | $r_s$ | $p$ | $r_s$ | $p$ |
| GBR-GRO |  |  | 0.306 | < 2.2e-16 |
| DEN-NOR | 0.147 | < 2.2e-16 | 0.364 | < 2.2e-16 |
| BEL-MAL | 0.174 | < 2.2e-16 | 0.389 | < 2.2e-16 |
| SWE-FIS | 0.137 | < 2.2e-16 | 0.332 | < 2.2e-16 |
| GER-RUE | 0.251 | < 2.2e-16 | 0.284 | < 2.2e-16 |
| FIN-HEL | 0.308 | < 2.2e-16 | 0.182 | < 2.2e-16 |
| SWE-BOL | 0.319 | < 2.2e-16 | 0.166 | < 2.2e-16 |
| FIN-KIV | 0.299 | < 2.2e-16 | 0.136 | < 2.2e-16 |
| SWE-BYN | 0.155 | < 2.2e-16 | -0.105 | < 1e-10 |
| SWE-KIR | 0.259 | < 2.2e-16 | 0.034 | 0.030 |
| FIN-KAR | 0.290 | < 2.2e-16 | -0.085 | < 1e-6 |
| FIN-PUL | 0.330 | < 2.2e-16 | 0.188 | < 2.2e-16 |
| FIN-RYT | 0.223 | < 2.2e-16 | -0.038 | 0.015 |

### Impact of introgression on population genetic analyses

Introgression affects population genetic summary statistics, such as nucleotide diversity (*π*), absolute divergence (*d_xy_*), and measure of population differentiation (*F_ST_*), and may seriously mislead inferences based on them. We found that the level of introgression was negatively correlated with *F_ST_* and *d_xy_* between the hybrid (NBS) and the minor (WL) parental population (GBR-GRO), but positively correlated when compared to the major (EL) parental population (RUS-BOL; Fig. 7a-b, Supplementary Figure S3). On the other hand, *π* in the hybrid (NBS) population was positively correlated with the level of introgression (Fig. 7c).

**Figure 7.**
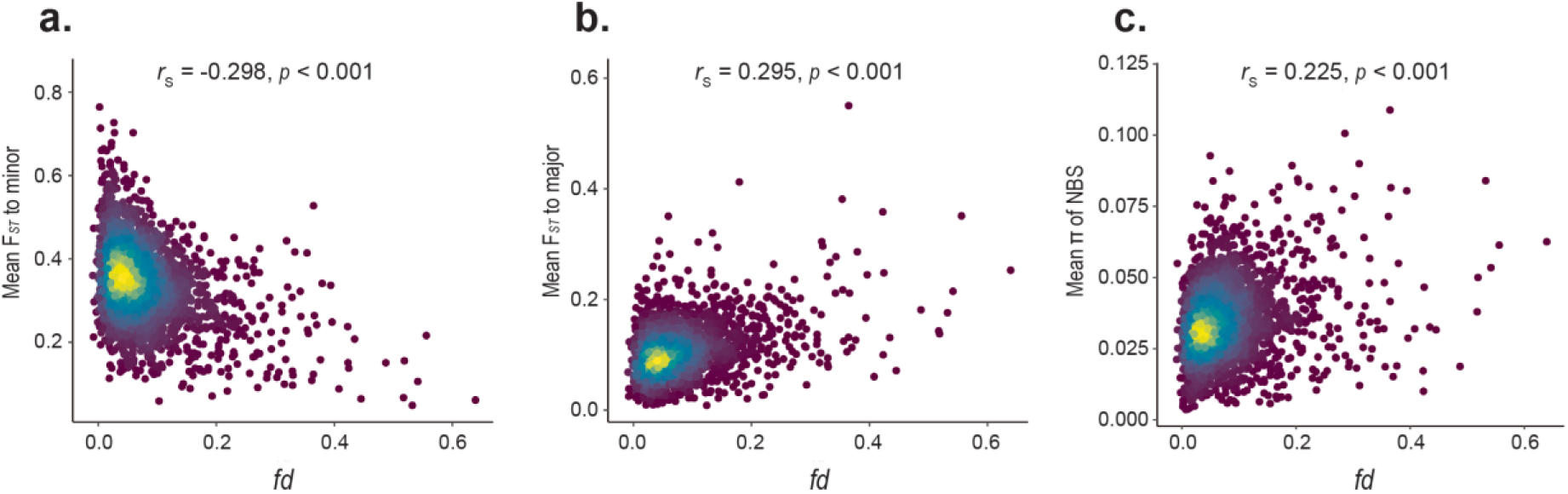
Effect of introgression on estimates of population differentiation (*F_ST_*) and nucleotide diversity (*π*) in the three northern Baltic Sea (NBS) populations. Correlation between mean estimates of admixture proportions (*fd*) and: **a.** *F_ST_* to minor (GBR-GRO) and **b.** major (RUS-BOL) parental population, and **c.** *π* of the NBS populations. Colours indicate density of points from low (dark blue) to high (yellow).

*F_ST_* is probably the most commonly used summary statistic in the identification of local adaptation (Beaumont *et al.* 2005; Narum and Hess 2011; Bierne *et al.* 2013). We studied how such analyses are affected by introgression when it has happened 1) in the target population being tested (often small *N_e_*); 2) in the reference population (often larger *N_e_*) being tested against, or 3) in both the test and reference population. For comparison, we analyzed two non-admixed populations. We specifically focused on the origin of outlier SNPs, that is, whether they were classified as WL- or EL-origin in NBS and the admixed target (SWE-BYN; Methods and Supplementary Table S8). The challenge of the analyses was that the correct outliers could not be distinguished. We therefore assumed that introgression from the marine WL is not beneficial in adaptation to pond environments in the EL dominated genetic background, and WL-origin alleles should not be overrepresented among the outliers.

In the first scenario, we compared admixed SWE-BYN with non-admixed RUS-BOL and found that 15.9% of the *F_ST_* outliers were WL-shared variants, while only 4.7% of the genome-wide variants were classified as such (Supplementary Table S8). Moreover, both the cut-off threshold of *F_ST_* (0.837 for *p* < 0.05) and genome-wide mean *F_ST_* (0.3 ± 0.27) were the highest among the four pairwise comparisons performed (Table 2), suggesting that introgression has affected the analysis. In the second scenario, a comparison of non-admixed FIN-RYT and admixed NBS, 2.1% of the outliers were classified as WL-origin, and of those 32% were identified as positively selected WL-NBS shared variants. In the third scenario, we compared admixed SWE-BYN and NBS, and found 4.8% of the outliers to be WL-shared variants, and 47.7% of these classified as positively selected WL-NBS shared variants in NBS. However, 79.9% of the outliers resulting from WL-origin variants showed higher VAF in SWE-BYN than in NBS. It is notable that the number of outliers resulting from WL-origin variants was significantly lower in comparison to an admixed (3,206 for SWE-BYN vs. NBS) than to a non-admixed reference population (7,878 for SWE-BYN vs. RUS-BOL).

**Table 2.** *F_ST_* outlier test of four pairwise comparisons.

| Focal | Reference | Outliers resulting from WL-shared variants | | | mean $F_{ST} \pm$<br>SD | $F_{ST}$ threshold<br>($p=0.05$) |
| --- | --- | --- | --- | --- | --- | --- |
| | | $n$ | Proportion of<br>all outliers | Positively<br>selected in<br>BS | | |
| **SWE-BYN** | RUS-BOL | 7878 | 0.159 | | $0.30 \pm 0.27$ | 0.837 |
| FIN-RYT | **NBS** | 1371 | 0.021 | 439 | $0.26 \pm 0.24$ | 0.733 |
| **SWE-BYN** | **NBS** | 3206 | 0.048 | 307 | $0.25 \pm 0.23$ | 0.717 |
| FIN-RYT | RUS-BOL | 0 | 0 | | $0.25 \pm 0.24$ | 0.736 |

NOTE.—NBS: the combined admixed northern Baltic Sea populations (FIN-HEL, SWE-BOL, FIN-KIV), $n$ : number of outliers, SD: standard deviation. Population label in bold letters refers to an admixed population.

### Impact of introgression on phylogenomic analysis

To test how introgression affects phylogenomic analyses, we reconstructed phylogenetic trees for representative individuals from the 14 populations using maximum likelihood (ML). In total, 997 ML trees were obtained from 1,000 randomly selected non-overlapping 100kb-regions. We recorded the placement of the individuals from the seven admixed populations and classified the trees as WL branch-off, EL branch-off, or unresolved. Excluding unresolved trees, the northern Baltic Sea individuals grouped with WL in 10.2–12.7% of cases, while those from the German coast were classified as WL branch-off in 24.8% of cases (Fig. 8b, Supplementary Table S10). A similar pattern was seen in the admixed WL populations, and the most eastern SWE-FIS was classified as EL branch-off in 38.3% of cases, whereas for individuals from the North Sea, the proportion was 13.4–13.6% (Fig. 8b, Supplementary Table S10). When all the 1,000 loci were concatenated together, the single best ML had monophyletic groups of WL and EL+Baltic Sea, both with 100% bootstrap support (Fig. 8a). The branching order of admixed individuals followed their geographic origin and the amount of foreign ancestry from either WL or EL (Fig. 1 and Fig. 8c-d). The single-locus trees showed clearly lower congruence with the concatenated ML tree than the bootstrap analysis, the support for the two groupings falling to 43 and 41% (Fig. 8a).

**Figure 8.**
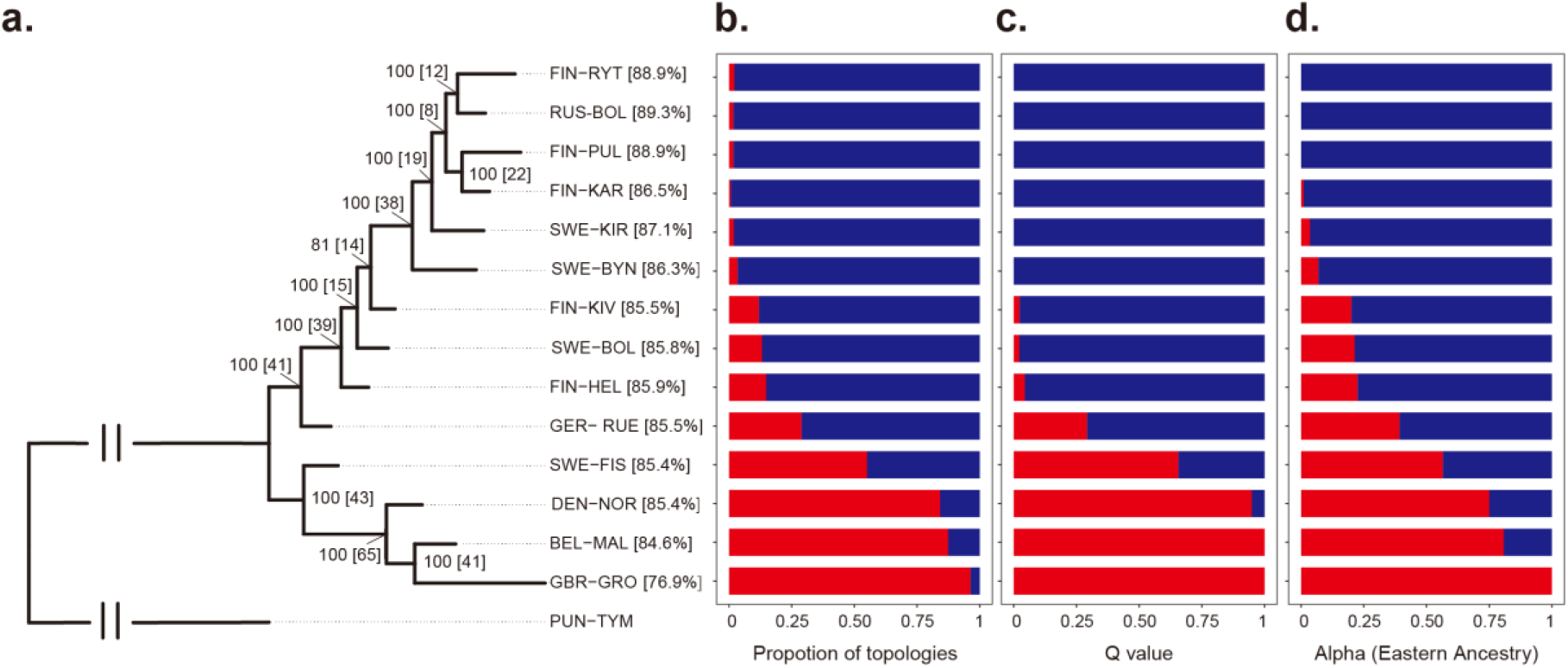
Phylogeny for representatives of each population shows intermediate placement of admixed populations. **a.** The tree shows the ML solution for the concatenated data with support values from 100 bootstrap replicates and from congruent single-locus trees (in brackets) for each tree node. Proportions of resolved trees for taxon placement are given after the taxon label. Red and blue show **b.** the proportion of resolved trees supporting the taxon’s monophyly with WL or EL, and **c.** the ADMIXTURE Q values (K=2) for the same individuals. **d.** Red and blue show the eastern (α) and western (1-α) ancestry for the population as estimated with *f4-ratio* test.

## Discussion

In the present study, we investigated the mechanisms and impact of genetic introgression using two divergent stickleback lineages (Fig. 1). Our inferences based on genetic variation from genome-wide autosomal SNPs confirm the earlier mitochondrial DNA based inference (Teacher *et al.* 2011) that the Danish straits constitute a contact zone between the Western and Eastern European nine-spined stickleback lineages. However, our results show that the hybrid zone actually extends from the southern and eastern North Sea to the northern Baltic Sea, and that WL introgression has penetrated also into pond populations currently isolated from the Baltic Sea (Fig. 2, Fig. 4c). We observed a gradient in foreign ancestry decreasing with increasing distance from the Danish straits, and demographic analyses revealed that the introgression may have started 10,000 ya (Fig. 3), soon after the reconnection of the North and Baltic Seas in the Mountain Billingen area (Schwarzer *et al.* 2008). Admixed populations from the Baltic Sea showed bursts in *N_e_* starting around 5,000 ya, close to the opening of the Danish straits 7,000 ya (Schwarzer *et al.* 2008). Signals of earlier demographic events may be hidden by the extensive levels of introgression observed, but a deep bottleneck was seen in the non-introgressed FIN-KAR around 12,000–16,000 ya (Fig. 3). This bottleneck may be related to the invasion of the Baltic Sea area from the eastern White Sea area after the last glacial period (Guo *et al.* 2019). We found several landlocked pond populations with low levels of WL admixture in the areas that used to be part of the Baltic Sea before the land uplift following the last glaciation isolated them from the sea (Mobley *et al.* 2011). The fact that WL introgression has penetrated these ponds confirms that the introgression from WL to EL is not only of recent origin, but must have occurred thousands of years ago. Despite a rapid increase in *N_e_*, we did not find evidence for the WL population GBR-GRO being admixed with the EL. Instead, the population may have experienced some level of admixture with *P. laevis* that is currently distributed in mainland Europe (Guo *et al.* 2019) or had a secondary contact with its surrogate marine or another distantly related population.

Recent studies suggest that introgression may be an important source of genetic variation fueling adaptive evolution and evidence for adaptive introgression has been found in a diverse array of taxa (Huerta-Sanchez *et al.* 2014; Sankararaman *et al.* 2014; Racimo *et al.* 2015, 2017; Dannemann *et al.* 2017; Richards and Martin 2017; Oziolor *et al.* 2019; Enciso-Romero *et al.* 2017; Suarez-Gonzalez *et al.* 2018; Hedrick 2013). Our results add to this evidence by identifying a set of genes, possibly related to adaptation to the marine environment, with high-frequency WL variants in EL genetic backgrounds that show footprints of selection (Supplementary Tables S7). *PIK3R6* has been shown to be differentially expressed between marine and freshwater nine-spined sticklebacks (Wang *et al.* 2020), and in response to salinity acclimation between marine and freshwater ecotypes in the three-spined stickleback (Gibbons *et al.* 2017). Moreover, *PIK3R6* is involved in the *Phosphoinositide 3-kinase* (*PI3K*) signaling pathway participating in hypoxia adaptation (reviewed in Zhu *et al*. 2013). Of the other AI genes, *PLEKHG3* was found to be a *PI3K*-regulated Rho guanine nucleotide exchange factor (Nguyen *et al.* 2016), involved in maintaining cell polarity and directional motility (Nguyen *et al.* 2016), and possibly important in response to infections (Van Keymeulen *et al.* 2006, Muthuswamy and Xue 2012) in brackish water environments. Interestingly, we also found evidence for adaptive introgression in *ZP4* that plays an important role in sperm-egg interactions during fertilization (Wassarman *et al.* 2001). The structure of the chorion, the coat of the egg cell, is an important adaptive feature for fish in brackish water conditions of the Baltic Sea (Lønning and Solemdal, 1972; Nissling *et al.* 2002). Although further work is needed to identify the actual targets of selection and biological functions of the candidate genes, it seems reasonable that the adaptively introgressed WL variants could have facilitated acclimatization to the brackish water environment of the Baltic Sea.

The influence of natural selection can be seen on the levels of introgression among different categories of genomic elements. We found WL ancestry in the admixed Baltic Sea populations to be significantly enriched in promoter regions and depleted in constrained elements (Supplementary Tables S5). This suggests that introgression is more likely to induce regulatory changes in gene expression than changes in protein-coding genes (Gittelman *et al.* 2016; Dannemann *et al.* 2017; McCoy *et al.* 2017; but see: Petr *et al.* 2019; Silvert *et al.* 2019), and that these changes happen at a very local scale, leaving central regulatory processes and developmental enhancers (Visel *et al*. 2008) less affected. While the different distributions of foreign ancestry in EL and WL populations may be explained by the lower overall levels of introgression in WL populations, also the environment and population demographic histories of the two introgression directions are very different. The invasion of EL from the White Sea to the Baltic Sea area was likely associated with a population bottleneck as well as a shift from saline to freshwater and then back to brackish water conditions. The WL introgression to the Baltic Sea populations may thus have been advantageous and facilitated EL fish to re-adapt to a marine-like (brackish water) environment. Introgression in the opposite direction may not have given similar advantages.

Recombination rate variation is known to play a key role in shaping the genomic landscape of introgression (e.g. Kim *et al.* 2018; Martin *et al.* 2019). Unlike previous studies (e.g. Schumer *et al.* 2018; Martin *et al.* 2019; Edelman *et al.* 2019; but see: Pool 2015), we found that the admixture proportions were mostly negatively correlated with recombination rates (Supplementary Table S9). The opposing results could be understood if genomic incompatibilities are the dominant force shaping the genomic landscape of introgression in the studies reporting positive correlations, but not in studies reporting negative correlations. In fact, the level of introgression can be negatively correlated with recombination rate when the deleterious variants are recessive, and the patterns are expected to be even stronger when the donor population has a large *N_e_* compared to the recipient population (Kim *et al.* 2018). Given that no hybrid incompatibility exists between WL and EL populations (Natri *et al*. 2019) and the historical population sizes appear to have been highly uneven, the negative correlation we observed may not be unexpected.

However, a negative correlation between *fd* and recombination rate may also result from selection favouring specific variants derived from the major parental population. We found the Baltic Sea polymorphism shared with WL to be largely neutral, and among the few positively selected sites (5,305), selection was more frequently acting on native EL-shared than on WL-shared genetic variants (90.23% vs. 9.77%; Fig. 5). If the EL origin variants have a small selective advantage, the selection will be most efficient in high recombination regions, driving a negative correlation between introgressed WL variation and recombination rate. Interestingly, if local adaptation plays a role, we should see its impact also in population differentiation, measured as *F_ST_*. Whereas linked selection can produce negative correlation between *F_ST_* and recombination rate through decrease in *N_e_* and increase in drift in low recombination regions, selection is more efficient in high recombination regions and produces positive correlation between *F_ST_* and recombination rate under divergent adaptation. This correlation would be strengthened if local adaptations are common and the targets for selection are enriched in high recombination regions (Keinan and Reich 2010). In line with that, both *F_ST_* (Table 1) and the density of genes (Varadharajan *et al.* 2019) are mostly positively correlated with recombination rate. Taken together, the results suggest that the two stickleback lineages have been under divergent selection on numerous loci and the selection on those loci has been strong enough to resist both the impact of drift and introgression.

The alternative approaches to quantify WL introgression to the Baltic Sea populations measured different signals and produced slightly contrasting results (Fig. 2, Fig. 8). Of the discrepancies between different approaches, the most interesting is the one between the ancestry proportions estimated with the *f_4_-ratio* test and the polymorphisms shared between the Baltic Sea hybrid populations and the two ancestral lineages. While nearly 80% of the hybrid population genomes are inferred to be of EL origin (77.4–79.9% in *f_4_-ratio’s* direct estimation of EL), they share only 16.3% of their polymorphism with EL and as much as 11.9% with WL. This can be explained by the bottleneck in the EL lineage colonizing the Baltic Sea, and the loss of much of the ancestral variation there. Interestingly, 90.2% of the positively selected ancestral variants are of EL origin, suggesting that, although much of the variation was lost, the important adaptive variation was retained through the bottleneck in the colonizing lineage. In contrast, a high proportion (99.84%) of the WL-origin polymorphism appeared to be selectively neutral, confirming the limited role of WL introgression in adaptation of the Baltic Sea sticklebacks.

We discovered that *F_ST_* and *d_xy_* can be both positively or negatively correlated with levels of introgression, depending on which parental population they are compared with (Fig. 7a-b, Supplementary Figure S3). On the other hand, *π* was always positively correlated with levels of introgression. As such, these results are perfectly expected and confirm that gene flow introduces new variation and makes populations more similar to the donor, but more differentiated from the other parental population. More important are the consequences of this, i.e. that unaccounted introgression may seriously alter local estimates of *π*, *F_ST_* and *d_xy_*. Of these, *F_ST_* has been widely used in identification of loci putatively involved in local adaptation (Hoban *et al.* 2016; Lotterhos and Whitlock, 2014; Matthey-Doret, 2019).

Although the difficulty of false-positive outliers and the effect of admixture are recognized (Lotterhos and Whitlock 2014; Hoban *et al.* 2016; Narum and Hess 2011; Nachman and Payseur 2012), the impact of introgression on our ability to discern local adaptation from outlier analyses is not well understood (but see Cruickshank and Hahn 2014; Hey 2010). By investigating *F_ST_* patterns of admixed populations, we demonstrated that especially introgression in the focal population can heavily affect the results. In a comparison of an admixed pond population to a non-admixed reference, we found outliers to be WL-shared variants three times more often than expected from the genome-wide estimation of ancestry proportions (Table 2). While many of the outliers are likely false inferences, introgression also increases the variance of *F_ST_* estimates and makes the detection of the true signal difficult. In a comparison to an admixed reference, 79.9% of the WL-NBS shared variants had a higher VAF in the admixed pond population: these WL-origin variants may have had adaptive value in the pond, but they may as well be explained by the small *N_e_* and the demographic history. On the other hand, if introgressed variants have been driven to high frequencies independently, a comparison of two admixed populations may underestimate the local adaptation. In our analyses, the impact of introgression in the reference population only seemed small.

If introgression has a general tendency to lead to overestimation of local adaptation, it may be instructive to note that stickleback fishes have spearheaded the study of local adaptation: numerous investigations have been conducted to understand how colonization of freshwater habitats by marine ancestors has molded the genetic architecture of freshwater populations (e.g. Hohenlohe *et al.* 2010; Jones *et al.* 2012; Fang *et al.* 2019). However, few – if any – of these studies have considered the possible effects of historical gene flow from a divergent lineage or species in both focal (freshwater, small *N_e_*) and reference (marine, large *N_e_*) populations, and the possibility of biases in *F_ST_* based inference should be re-examined. Besides the test statistics, unaccounted introgression may affect the studies of parallel local adaptation. Isolated populations have generally been regarded as independent natural replicates (e.g. Jones *et al.* 2012), and unaccounted introgression may lead to heavily biased results and interpretations. Introgression increases the levels of standing genetic variation in admixed populations, providing more targets for selection and thereby differentiating the patterns of parallelism among admixed populations from those in non-admixed ones. Observing non-parallel patterns of local adaptation in studies where one regional set of populations has experienced introgression or secondary contact, and others have not, should not come as a surprise. Finally, introgression can also bias others tests requiring a neutral baseline of differentiation such as quantitative trait analyses (e.g. Leinonen *et al.* 2013).

Despite no single tree correctly representing phylogenetic relationships across the genome, phylogenetic trees are commonly inferred for closely related individuals (Martin *et al.* 2019; Stankowski *et al.* 2019). Our analyses provide a genome-wide view on the effects of introgression on tree topologies and illustrate that the probability of a sample being placed in the WL or EL lineage correlates with the amount of WL introgression in the population. It is also noteworthy to remind that the concatenation of large genomic data can hide the topological heterogeneity and give seemingly strong support for a single tree (Salichos and Rokas 2013). We obtained 100% support for all but one node in a bootstrap analysis of the full concatenated data while the phylogenetic congruence support varied from 8% to 65% across the nodes of the tree (Fig. 8), clearly better reflecting the underlying signal. We also observed the specific biases of trees inferred from admixed samples such as intermediate placement of admixed samples and shorter distances between the admixed sample and its sources than between the two sources (Kopelman *et al.* 2013).

In conclusion, our comprehensive study of genetic introgression between two divergent sticklebacks lineages shows that introgression is geographically widespread, and to some extent, has contributed to local adaptation in the recipient populations in the Baltic Sea area. Although the observed negative correlation between levels of introgression and recombination rate is a stark contrast to findings in most earlier studies (e.g. Schumer *et al.* 2018, Martin *et al.* 2019; Edelman *et al.* 2019; Stankowski *et al.* 2019), it is consistent with the observed positive correlation between population differentiation and recombination rate, as well as with our analyses of positive selection. The results demonstrate that genetic recombination and selection on ancestral standing variation can shape the genomic landscape of both introgression and differentiation in unexpected ways, though further work is still required to understand how general our findings are. Finally, our work has important practical implications for studies of local adaptation. With a growing number of studies showing widespread introgression among divergent lineages, it may be advisable to analyze possible effects of admixture before committing to analyses of local adaptation. As demonstrated by our results, unaccounted admixture can profoundly affect the interpretation of commonly used population genetic summary statistics.

## Methods

### Sample collection

The samples used in this study were collected in accordance with the national legislation of the countries concerned. A total of 289 nine-spined stickleback individuals (15–27 per population) were sampled from two earlier identified evolutionary lineages, the EL (ten populations) and the WL (four populations). The lineage assignment of populations was based on information from earlier studies (Teacher et al. 2011, Guo et al. 2019), and confirmed with data from this study (see Results). The samples were collected during the local breeding seasons via seine nets and minnow traps. After anesthetizing the fish with an overdose of MS-222, whole fish or fin clips were preserved in 95% ethanol and stored at - 80°C until DNA extraction. In addition, one *P. tymensis* individual serving as an outgroup was collected from Hokkaido, Japan (43°49’40’’N,145°5’10”E). The sampling sites are shown in Fig. 1, more detailed information, including sampling site coordinates and dates, sample sizes, population codes, and species names are given in Supplementary Table S1.

### DNA extraction and whole-genome resequencing

Extractions of genomic DNA were conducted following the standard phenol-chloroform method (Sambrook and Russell 2006) from alcohol-preserved fin clips. DNA libraries with an insert size of 300–350 bp were constructed, and 150-bp paired-end reads were generated using an Illumina HiSeq 2500/4000 instrument. Library preparations and sequencing were carried out at the Beijing Genomics Institute (Hong Kong, China) and the DNA Sequencing and Genomics Laboratory, University of Helsinki (Helsinki, Finland).

### Sequence alignment and variant calling

The reads were mapped to the nine-spined stickleback reference genome (Varadharajan *et al.* 2019) using the Burrows-Wheeler Aligner v.0.7.17 (BWA-MEM algorithm, Li 2013) and its default parameters. Duplicate reads were marked with samtools v.1.7 (Li *et al.* 2009) and variant calling was performed with the Genome Analysis Toolkit (GATK) v.4.0.1.2 (McKenna *et al.* 2010) following the GATK Best Practices workflows. In more detail, RealignerTargetCreator and IndelRealigner tools were applied to detect misalignments and realign reads around indels. The HaplotypeCaller was used to call variants for each individual, parameters set as -stand_emit_conf 3, -stand_call_cof 10, -GQB (10,50), variant index type linear and variant index parameter 128000. Then GenotypeGVCFs algorithms were then used to jointly call the variants for all the samples using default parameters. Interspecific variants were removed and binary SNPs were extracted with bcftools v.1.7 (Li *et al.* 2009), excluding sites located within identified repetitive sequences (Varadharajan *et al*. 2019). Sites showing an extremely low (< 5x) or high average coverage (> 25x) and quality score < 30 were filtered out using vcftools v.0.1.5 (Danecek *et al.* 2011). For subsequent filterings of dataset used in different analyses, see Supplementary Table S2 for details.

### Analysis of population genetic structure

The approximate population structure among all study samples was estimated using PCA within the PLINK toolset v.1.90 (Purcell *et al.* 2007) and ancestry estimation within ADMIXTURE v.1.3.0 (Alexander *et al.* 2009). In the latter, the number of ancestral populations (*K*) was set to 2 to represent WL and EL. In both PCA and ADMIXTURE analysis, the distance between two neighboring SNPs was restricted to be 10 kb to control for the effect of linkage disequilibrium.

### Test and quantification of genomic introgression

Patterson’s D statistic (Patterson *et al.* 2012) was used to assess if certain populations share more derived alleles with a donor population than is expected by chance. Using the typical topology of ((P1, P2), P3; O),

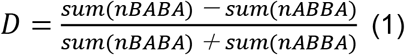

and an excess of BABA genealogies indicates gene flow between P1 and P3. We used *P. tymensis* always as an outgroup (O), and performed two distinct tests:

1. To assess EL introgression to WL populations, we used RUS-BOL, FIN-RYT, and FIN-PUL as P3; and GBR-GRO and BEL-MAL as P2 and P1, or P1 and P2, respectively.
2. To assess WL introgression in all other populations, we used GBR-GRO as P3, RUS-BOL, FIN-RYT, and FIN-PUL as P2 and the rest of the populations (except BEL-MAL) as P1.

For the populations yielding significant (i.e. *Z* ≥ 3) estimates of the D-statistic, the f_4_-ratio test (Reich *et al*. 2009) was applied to quantify the amount of foreign ancestry. Following Petr *et al.* (2019), the direct estimation of EL ancestry (α_EL_) was estimated as:

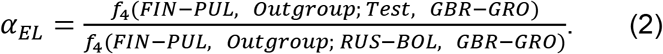

Assuming no admixture from other groups, the WL ancestry is then 1 - α_EL_.

Direct estimation of WL ancestry (α_WL_) was estimated as:

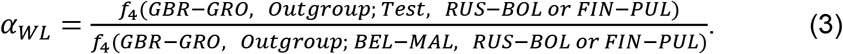

We caution that this may underestimate the true proportion of WL ancestry as BEL-MAL is admixed with EL. *D*-statistics and *f_4_-ratio* tests were performed using ADMIXTOOLS v.5.1 (qpDstat v.755, qpF4ratio v.320; Patterson *et al.* 2012).

To quantify the amount of WL and EL ancestry in different genomic features, the genome was binned to intergenic, CDS, constrained elements, introns and promoters based on the genome annotation (Varadharajan *et al.* 2019). Promoter regions were defined as 1kb-stretches upstream of the gene start, and locations of constrained elements were lifted over from the three-spined stickleback genome annotation (Ensembl release ver. 95) using Crossmap v.0.3.3 (Zhao *et al.* 2014). A significance test was applied to assess whether EL and WL ancestry were significantly enriched or depleted in any of the genomic features. Following Petr *et al.* (2019), the *α* value of a given annotation category was resampled 10,000 times from a normal distribution centered on the *α* with a standard deviation equal to the standard error given by ADMIXTOOLS. An empirical *p*-value was then calculated for the estimated *α* for each genomic feature to test the hypothesis that the ancestry proportion does not differ from that of the intergenic regions.

### Reconstruction of demographic history

SMC++ (v.1.15.3; Terhorst *et al.* 2017) was used to reconstruct the demographic histories of the study populations because of its ability to utilize unphased data from multiple individuals. The individual with the highest sequencing coverage (in most cases 20X) was set as ‘distinguished individual’, and mutation rate of 1.42×10^-8^ per site per generation (Guo *et al.* 2013) and generation time of 2 years (De Faveri *et al.* 2014) were assumed. To focus on the last glacial period (15-110 kya) and opening of the connection between the Baltic Sea and the North Sea (~10 kya; Schwarzer *et al.* 2008), the time interval of inference was limited to 1,000–1×10^6^ generations ago. To avoid overfitting, we set --regularization-penalty 5 --ftol 0.01 --folds 4.

### Quantification of introgression across the genome (*fd*)

The modified *D*-statistic, *fd* (Martin *et al.* 2015), was used to quantify introgression at finer genomic scales. We used a fixed window size of 100-kb with a 20-kb step size using the scripts from Martin *et al.* (2015) and estimated two-tailed *p*-values from the *Z*-transformed *fd* values using the standard normal distribution and corrected for multiple testing with the Benjamini-Hochberg false discovery rate (FDR) method (Benjamini and Hochberg 1995). Windows with positive *D* and *fd* values with a number of informative sites ≥ 100 and FDR value ≤ 0.05 were retained as outlier loci (see below). To minimize population-specific effects (such as inversions) in the reference population, three non-admixed populations (FIN-PUL, FIN-RYT, RUS-BOL) were used alternately as the EL source for the three NBS populations; GBR-GRO was always used as the WL source, giving nine pairwise comparisons. Intersections of all outlier loci among the nine pairwise computations were considered as putative introgression-enriched regions, merging overlapping regions into one.

Read coverage and called genotypes were used to classify the introgression events and to rule out false inferences e.g. due to structural variants. Introgressed regions were defined as duplicates or deletions if the average read coverage was above 20 or below 1, respectively, and there was an excess of heterozygous or missing genotypes across the regions shared between GBR-GRO and the three Baltic Sea populations. The length of the introgressed segments (*m*, see below) was defined as the distance between the first and last SNP within each region.

Introgression was distinguished from ILS using a test based on recombination rate. The expected length of a shared ancestral sequence due to ILS can be calculated as *L*=1/(*r*t*), where *r* is the local recombination rate (scaled as per base-pair per generation) and *t* is the number of generations since divergence. Following Huerta-Sánchez *et al.* (2014), we assumed an exponential distribution for the length of the introgressed tracks with the probability of observing a shared track of length ≥ *m* (in base pairs) being *exp*(*-m/L*). Conditional on observing the WL nucleotide at position *j*, the expected length is the sum of two exponentially distributed random variables with expected lengths *L*, which follows a Gamma distribution with shape parameter 2 and a rate parameter lambda = *1/L*. Thus, the probability of each of the introgressed segments was computed as the probability of observing a fragment of at least a given length, *m*, as 1-Gamma(*m*, shape=2, rate=*1/L*). This probability was estimated separately for each of the introgressed segments. The *t* was set to be 215 000, based on 0.43 Mya divergence between WL and EL and generation time of two years, and the mean local recombination rate *r* was obtained from Varadharajan *et al.* (2019) binned to the size of the candidate regions.

### Footprints of selection in Baltic Sea populations

Footprints of selection on introgressed variants were searched using the *U* and *Q*95 tests following Racimo *et al.* (2017). Both tests are based on the variant allele frequencies (VAF) and focus, respectively, on the number of SNPs shared with donor population at high frequency in a focal population but at low frequency in a reference population, and the 95% quantile of the frequency of the SNPs that are shared with a donor population and at low frequency in a reference population. The VAF was estimated separately for WL (GBR-GRO), EL (FIN-RYT, FIN-PUL, RUS-BOL), and NBS (FIN-HEL, SWE-BOL, FIN-KIV) populations, and the tests were performed using 100-kb windows with a 20-kb step and discarding positions with more than 25% missing data. We first calculated the *U*20_EL, NBS, WL_(0.01, 0.2, 1) to count the SNPs that are at < 1% frequency in the combined EL reference population, at ≥ 20% frequency in the combined NBS population, and fixed (100% frequency) in the WL population. We then calculated the *Q*9_5EL_, NBS, WL(0.01, 0.2, 1) to obtain the 95% quantile VAF of these SNPs in NBS. The intersection of the top 1% regions of the *U*20 and *Q*95 tests and the candidate regions from the *fd* test were considered as putative adaptive introgression (AI) regions. Within each AI region, *F_ST_*, *d_xy_*, and *π* were calculated for 10-kb sized windows with a 5-kb step using the scripts from Martin *et al.* (2015), sequencing coverage, SNP information from *U* and *Q*95 test results, and variant allele frequencies (minor allele frequency [maf] ≥ 0.05) were used to identify candidates for possible adaptive evolution among the gene annotations (Varadharajan *et al.* 2019).

The VAFs were also used to assess the relative contributions of introgression (WL origin) and ancestral standing genetic variation (EL origin). VAFs were estimated for all variants in WL (GBR-GRO), EL (RUS-BOL), and NBS (FIN-HEL, SWE-BOL, FIN-KIV) populations, discarding sites with more than 25% missing data or VAF_NBS_ = 0. We defined the variants in the Baltic Sea populations to be of NBS-EL or WL-NBS shared based on their presence/absence and absence/presence in RUS-BOL and GBR-GRO, respectively. Based on these two sets of SNPs, we then classified the variants in the Baltic Sea populations as follows. (1) WL-NBS shared SNPs (VAF_WL_>0 and VAF_EL_=0) were defined as *WL-origin;* (2) WL-NBS shared SNPs with VAF_NBS_>0.84, which is above the 99% quantile of the *Q*95 test scores, were defined as *selection on WL-NBS shared variants* (as a proxy of adaptive introgression), the rest being neutral (as a proxy of neutral introgression); and (3) EL-NBS shared SNPs with VAF_NBS_>0.88, which is above the 99% quantile of the SNP category (VAF_NBS_[VAF_EL_>0 and VAF_WL_=0]), were defined as *selection on ancestral variants*.

### Impact of introgression on population genetics analyses

We examined the covariation of admixture proportions (*fd*) with several population genetic statistics: nucleotide diversity (*π*), absolute divergence (*d_xy_*), and measure of population differentiation (*F_ST_*). The statistics were computed genome-wide in 100-kb non-overlapping windows. The mean recombination rate was estimated from the linkage map (Varadharajan *et al.* 2019) for 10-kb windows and binned into 100-kb non-overlapping windows.

To evaluate the effect of introgression on *F_ST_*-based outlier detection, the per-site *F_ST_* was computed using vcftools v.0.1.5 (Danecek *et al.* 2011). Non-admixed FIN-RYT and admixed SWE-BYN pond populations were used as test populations, and non-admixed RUS-BOL and the three admixed Baltic Sea populations (NBS; grouped together) as the reference populations, requiring maf > 0.05 to reduce background noise. Negative *F_ST_* values were discarded and *p*-values were estimated from the *Z*-transformed *F_ST_* values using the standard normal distribution. *F_ST_* estimates with *p* ≤ 0.05 were regarded as outliers.

### Phylogenomic analysis

Phylogenetic relationships were estimated with RAxML v.8.2.9 (Stamatakis *et al.* 2014), selecting randomly one individual from each population and using *P. tymensis* as the outgroup. To capture the effects of ILS and variation in evolutionary history across sites, 1,000 non-overlapping regions, each 100-kb in size, were randomly selected. The ‘GTRGAMMA’ model was used and *--asc-corr=lewis* was applied to correct for ascertainment bias in the SNP data. The per-locus maximum likelihood (ML) trees were classified as WL monophyletic, EL monophyletic, and paraphyletic using the *ape* package v.5.3 (Paradis *et al.* 2019) in R v.3.5.1. A consensus phylogeny was constructed using the concatenated sequence of the 1,000 loci and support values obtained from 100 bootstrap replicates, inferred using the same parameters.

## Data availability

Raw sequence data will be available at European Nucleotide Archive (ENA) (https://www.ebi.ac.uk/ena) under accession code PRJEB38005. Other relevant data are available from the corresponding author upon request.

## Code availability

All code and scripts for downstream analysis are available at https://github.com/XueyunF/Introgression-in-Ninespined-stickleback

## Supplementary information

Supplementary Information

Supplementary Table S3_S4

Reporting Summary

## Acknowledgments

We thank Victor Berger, Pär Byström, Lasse Fast Jensen, Jacquelin De Faveri, Gabor Herczeg, Tuomas Leinonen, Scott McCairns, Andrew McColl, Heini Natri, Takahito Shikano and, Joost Raeymaekers for help in providing samples, and Miinastiina Issakainen, Sami Karja, and Kirsi Kähkönen for help in the laboratory; Bohao Fang for help in data visualization. The advice and support from Antoine Fraimout, Baocheng Guo, Cui Wang, Paolo Momigliano, Pasi Rastas, Petri Kemppainen, Zitong Li, Mikko Kivikoski, Jarkko Salojärvi, Jack Beresford, Jacquelin De Faveri, Takahito Shikano, Simon Martin, and Martin Petr is gratefully acknowledged. Our research was supported by grants from the Academy Finland (# 129662, 134728 and 218343 to JM; # 322681 to AL), Helsinki Lifesciences Center (HiLife; to JM), and China Scholarship Council (# 201608520032 to XF). Computational resources provided by the CSC–IT Center for Science, Finland, are acknowledged with gratitude.

## Author contributions

A.L and J.M conceived the original idea, with significant later contributions from X.F. X.F and A.L analyzed the data. X.F took lead in writing the manuscript, with significant contributions from A.L and J.M.

## Ethics declarations

## References

1. Alexander DH, Novembre J, Lange K. Fast model-based estimation of ancestry in unrelated individuals. Genome Research, 2009, 19(9): 1655–1664.

2. Barton N, Bengtsson BO. The barrier to genetic exchange between hybridising populations. Heredity, 1986, 57(3): 357–376.

3. Bay RA, Taylor EB, Schluter D. Parallel introgression and selection on introduced alleles in a native species. Molecular Ecology, 2019, 28(11): 2802–2813.

4. Beaumont MA. Adaptation and speciation: what can F_ST_ tell us?. Trends in Ecology & Evolution, 2005, 20(8): 435–440.

5. Benjamini Y, Hochberg Y. Controlling the false discovery rate: a practical and powerful approach to multiple testing. Journal of the Royal statistical society: series B (Methodological), 1995, 57(1): 289–300.

6. Bierne N, Roze D, Welch JJ. Pervasive selection or is it…? Why are F_ST_ outliers sometimes so frequent?. Molecular Ecology, 2013, 22(8): 2061–2064.

7. Bomblies K, Lempe J, Epple P, et al. Autoimmune response as a mechanism for a Dobzhansky-Muller-type incompatibility syndrome in plants. PLoS Biology, 2007, 5(9).

8. Cruickshank TE and Hahn MW. Reanalysis suggests that genomic islands of speciation are due to reduced diversity, not reduced gene flow. Molecular Ecology, 2014, 23.13: 3133–3157.

9. Danecek P, Auton A, Abecasis G, et al. The variant call format and VCFtools. Bioinformatics, 2011, 27(15): 2156–2158.

10. Dannemann M, Prüfer K, Kelso J. Functional implications of Neandertal introgression in modern humans. Genome Biology, 2017, 18(1): 61.

11. DeFaveri J, Shikano T, Merilä J. Geographic variation in age structure and longevity in the nine-spined stickleback *(Pungitius pungitius)*. PLoS One, 2014, 9(7).

12. Edelman NB, Frandsen PB, Miyagi M, et al. Genomic architecture and introgression shape a butterfly radiation. Science, 2019, 366(6465): 594–599.

13. Enciso-Romero J, Pardo-Díaz C, Martin SH, et al. Evolution of novel mimicry rings facilitated by adaptive introgression in tropical butterflies. Molecular Ecology, 2017, 26(19): 5160–5172.

14. Fang B, Kemppainen P, Momigliano P, et al. Oceans apart: Heterogeneous patterns of parallel evolution in sticklebacks. Preprint at https://www.biorxiv.org/content/10.1101/826412v2 (2020).

15. Gibbons TC, Metzger DCH, Healy TM, et al. Gene expression plasticity in response to salinity acclimation in threespine stickleback ecotypes from different salinity habitats. Molecular Ecology, 2017, 26(10): 2711–2725.

16. Gittelman RM, Schraiber JG, Vernot B, et al. Archaic hominin admixture facilitated adaptation to out-of-Africa environments. Current Biology, 2016, 26(24): 3375–3382.

17. Guo B, Chain F JJ, Bornberg-Bauer E, et al. Genomic divergence between nine- and three-spined sticklebacks. BMC Genomics, 2013, 14(1): 756.

18. Guo B, Fang B, Shikano T, et al. A phylogenomic perspective on diversity, hybridization and evolutionary affinities in the stickleback genus *Pungitius*. Molecular Ecology, 2019, 28(17): 4046–4064.

19. Harris K, Nielsen R. The genetic cost of Neanderthal introgression. Genetics, 2016, 203(2): 881–891.

20. Hawks J. Introgression makes waves in inferred histories of effective population size. Human Biology, 2017, 89(1): 67–80.

21. Hedrick PW. Adaptive introgression in animals: examples and comparison to new mutation and standing variation as sources of adaptive variation. Molecular Ecology, 2013, 22(18): 4606–4618.

22. Henn BM, Botigué LR, Peischl S, et al. Distance from sub-Saharan Africa predicts mutational load in diverse human genomes. Proceedings of the National Academy of Sciences USA, 2016, 113(4): E440–E449.

23. Hey J. Isolation with migration models for more than two populations. Molecular Biology and Evolution, 2010, 27, 905–920.

24. Hoban S, Kelley JL, Lotterhos KE, et al. Finding the genomic basis of local adaptation: pitfalls, practical solutions, and future directions. American Naturalist, 2016, 188(4): 379–397.

25. Hohenlohe PA, Bassham S, Etter PD, et al. Population genomics of parallel adaptation in threespine stickleback using sequenced RAD tags. PLoS Genetics, 2010, 6(2).

26. Huerta-Sánchez E, Jin X, Bianba Z, et al. Altitude adaptation in Tibetans caused by introgression of Denisovan-like DNA. Nature, 2014, 512(7513): 194–197.

27. Jokinen H, Momigliano P, Merilä J. From ecology to genetics and back: the tale of two flounder species in the Baltic Sea. ICES Journal of Marine Science, 2019, 76(7): 2267–2275

28. Jones FC, Grabherr MG, Chan YF, et al. The genomic basis of adaptive evolution in threespine sticklebacks. Nature, 2012, 484(7392): 55–61.

29. Juric I, Aeschbacher S, Coop G. The strength of selection against Neanderthal introgression. PLoS Genetics, 2016, 12(11): e1006340.

30. Keinan A, Reich D. Human population differentiation is strongly correlated with local recombination rate. PLoS Genetics, 2010, 6(3).

31. Kim BY, Huber CD, Lohmueller KE. Deleterious variation shapes the genomic landscape of introgression. PLoS Genetics, 2018, 14(10): e1007741.

32. Kopelman NM, Stone L, Gascuel O, et al. The behavior of admixed populations in neighbor-joining inference of population trees. Biocomputing, 2013, 2013: 273–284.

33. Kuhlwilm M, Gronau I, Hubisz MJ, et al. Ancient gene flow from early modern humans into Eastern Neanderthals. Nature, 2016, 530(7591): 429–433.

34. Lamichhaney S, Han F, Webster MT, et al. Rapid hybrid speciation in Darwin’s finches. Science, 2018, 359(6372): 224–228.

35. Lee HY, Chou JY, Cheong L, et al. Incompatibility of nuclear and mitochondrial genomes causes hybrid sterility between two yeast species. Cell, 2008, 135(6): 1065–1073.

36. Leinonen T, McCairns RJS, O’Hara RB, et al. QST–FST comparisons: evolutionary and ecological insights from genomic heterogeneity. Nature Reviews Genetics, 2013, 14(3): 179–190.

37. Li, Heng, et al. The sequence alignment/map format and SAMtools. Bioinformatics, 2009, 25.16: 2078–2079.

38. Li H. Aligning sequence reads, clone sequences and assembly contigs with BWA-MEM. arXiv preprint arXiv:1303.3997, 2013.

39. Lotterhos KE, Whitlock MC. Evaluation of demographic history and neutral parameterization on the performance of F_ST_ outlier tests. Molecular Ecology, 2014, 23(9): 2178–2192.

40. Lønning S, and Solemdal P. 1972. The relation between thickness of chorion and specific gravity of eggs from Norwegian and Baltic flatfish populations. Fiskeridirektoratets Skrifter. Serie Havundersøkelse, 16: 77–88.

41. Malinsky M, Svardal H, Tyers AM, et al. Whole-genome sequences of Malawi cichlids reveal multiple radiations interconnected by gene flow. Nature Ecology & Evolution, 2018, 2(12): 1940–1955.

42. Marques DA, Meier JI, Seehausen O. A combinatorial view on speciation and adaptive radiation. Trends in Ecology & Evolution, 2019.

43. Martin SH, Davey JW, Jiggins CD. Evaluating the use of ABBA–BABA statistics to locate introgressed loci. Molecular Biology and Evolution, 2015, 32(1): 244–257.

44. Martin SH, Davey JW, Salazar C, et al. Recombination rate variation shapes barriers to introgression across butterfly genomes. PLoS Biology, 2019, 17(2): e2OO6288.

45. Martin SH, Jiggins CD. Interpreting the genomic landscape of introgression. Current Opinion in Genetics & Development, 2017, 47: 69–74.

46. Masly JP, Presgraves DC. High-resolution genome-wide dissection of the two rules of speciation in *Drosophila*. PLoS Biology, 2007, 5(9).

47. Matthey-Doret R, Whitlock MC. Background selection and F_ST_: consequences for detecting local adaptation. Molecular Ecology, 2019, 28(17): 3902–3914.

48. McCoy RC, Wakefield J, Akey JM. Impacts of Neanderthal-introgressed sequences on the landscape of human gene expression. Cell, 2017, 168(5): 916–927. e12.

49. McKenna A, Hanna M, Banks E, et al. The Genome Analysis Toolkit: a MapReduce framework for analyzing next-generation DNA sequencing data. Genome Research, 2010, 20(9): 1297–1303.

50. Mobley KB, Lussetti D, Johansson F, Englund G & Bokma F. (2011). Morphological and genetic divergence in Swedish postglacial stickleback *(Pungitius pungitius)* populations. BMC Evolutionary Biology, 11(1), 287.

51. Muthuswamy SK, Xue B (2012) Cell polarity as a regulator of cancer cell behavior plasticity. Annual Review of Cell and Developmental Biology, 2012, 28: 599–625.

52. Nachman MW, Payseur BA. Recombination rate variation and speciation: theoretical predictions and empirical results from rabbits and mice. Philosophical Transactions of the Royal Society B: Biological Sciences, 2012, 367(1587): 409–421.

53. Narum SR, Hess JE. Comparison of F_ST_ outlier tests for SNP loci under selection. Molecular Ecology Resources, 2011, 11: 184–194.

54. Natri HM, Merilä J, Shikano T. The evolution of sex determination associated with a chromosomal inversion. Nature Communications, 2019, 10(1): 1–13.

55. Nguyen TTT, Park WS, Park BO, et al. PLEKHG3 enhances polarized cell migration by activating actin filaments at the cell front. Proceedings of the National Academy of Sciences USA, 2016, 113(36): 10091–10096.

56. Nissling A, Westin L, and Hjerne O. 2002. Reproductive success in relation to salinity for three flatfish species, dab *(Limanda limanda)*, plaice *(Pleuronectes platessa)*, and flounder *(Pleuronectes flesus)*, in the brackish water Baltic Sea. ICES Journal of Marine Science, 59: 93–108.

57. Oziolor EM, Reid NM, Yair S, et al. Adaptive introgression enables evolutionary rescue from extreme environmental pollution. Science, 2019, 364(6439): 455–457.

58. Palumbi SR: Genetic divergence, reproductive isolation, and marine speciation. Annual Review of Ecology and Systematics, 1994, 25(1): 547–572.

59. Paradis E, Schliep K. ape 5.0: an environment for modern phylogenetics and evolutionary analyses in R. Bioinformatics, 2019, 35(3): 526–528.

60. Patterson N, Moorjani P, Luo Y, et al. Ancient admixture in human history. Genetics, 2012, 192(3): 1065–1093.

61. Peischl S, Excoffier L. Expansion load: recessive mutations and the role of standing genetic variation. Molecular Ecology, 2015, 24(9): 2084–2094.

62. Petr M, Pääbo S, Kelso J, et al. Limits of long-term selection against Neandertal introgression. Proceedings of the National Academy of Sciences USA, 2019, 116(5): 1639–1644.

63. Pool JE. The mosaic ancestry of the Drosophila genetic reference panel and the *D. melanogaster* reference genome reveals a network of epistatic fitness interactions. Molecular Biology and Evolution, 2015, 32(12): 3236–3251.

64. Purcell S, Neale B, Todd-Brown K, et al. PLINK: a tool set for whole-genome association and population-based linkage analyses. The American Journal of Human Genetics, 2007, 81(3): 559–575.

65. Racimo F, Marnetto D, Huerta-Sánchez E. Signatures of archaic adaptive introgression in present-day human populations. Molecular biology and evolution, 2017, 34(2): 296–317.

66. Racimo F, Sankararaman S, Nielsen R, et al. Evidence for archaic adaptive introgression in humans. Nature Reviews Genetics, 2015, 16(6): 359.

67. Reich D, Thangaraj K, Patterson N, et al. Reconstructing Indian population history. Nature, 2009, 461(7263): 489–494.

68. Richards EJ, Martin CH. Adaptive introgression from distant Caribbean islands contributed to the diversification of a microendemic adaptive radiation of trophic specialist pupfishes. PLoS Genetics, 2017, 13(8).

69. Salichos L, Rokas A. Inferring ancient divergences requires genes with strong phylogenetic signals. Nature, 2013, 497(7449): 327–331.

70. Sambrook J, Russell DW. (2006). Isolation of high-molecular-weight DNA from mammalian cells using formamide. In: Press CSHL (ed) Cold Spring Harb Protoc, 3rd edn. Cold Spring Harbor, NY.

71. Sankararaman S, Mallick S, Patterson N & Reich D. The combined landscape of Denisovan and Neanderthal ancestry in present-day humans. Current Biology, 2016, 26(9), 1241–1247.

72. Sankararaman S, Mallick S, Dannemann M, et al. The genomic landscape of Neanderthal ancestry in present-day humans. Nature, 2014, 507(7492): 354–357.

73. Schumer M, Xu C, Powell DL, et al. Natural selection interacts with recombination to shape the evolution of hybrid genomes. Science, 2018, 360(6389): 656–66O.

74. Schwarzer K, Ricklefs K, Bartholomä A, et al. Geological development of the North Sea and the Baltic Sea. Die Küste, 74 ICCE, 2008 (74): 1–17.

75. Silvert M, Quintana-Murci L, Rotival M. Impact and evolutionary determinants of Neanderthal introgression on transcriptional and post-transcriptional regulation. American Journal of Human Genetics, 2019, 104(6): 1241–1250.

76. Skoglund P, Ersmark E, Palkopoulou E, et al. Ancient wolf genome reveals an early divergence of domestic dog ancestors and admixture into high-latitude breeds. Current Biology, 2015, 25(11): 1515–1519.

77. Stamatakis A. *RAxML* version 8: a tool for phylogenetic analysis and post-analysis of large phylogenies. Bioinformatics, 2014, 30(9): 1312–1313.

78. Stankowski S, Chase M A, Fuiten A M, et al. Widespread selection and gene flow shape the genomic landscape during a radiation of monkeyflowers. PLoS Biology, 2019, 17(7): e3000391.

79. Suarez-Gonzalez A, Lexer C, Cronk QCB. Adaptive introgression: a plant perspective. Biology Letters, 2018, 14(3): 20170688.

80. Teacher AGF, Shikano T, Karjalainen ME, et al. Phylogeography and genetic structuring of European nine-spined sticklebacks *(Pungitius pungitius)—*mitochondrial DNA evidence. PLoS One, 2011, 6(5).

81. Terhorst J, Kamm JA, Song YS. Robust and scalable inference of population history from hundreds of unphased whole genomes. Nature Genetics, 2017, 49(2): 303.

82. Van Keymeulen A, et al. To stabilize neutrophil polarity, PIP3 and Cdc42 augment RhoA activity at the back as well as signals at the front. Journal of Cell Biology, 2006, 174(3): 437–445.

83. Varadharajan S, Rastas P, Löytynoja A, et al. A high-quality assembly of the nine-spined stickleback *(Pungitius pungitius)* genome. Genome Biology and Evolution, 2019, 11(11): 3291–3308.

84. Visel A, Prabhakar S, Akiyama JA, et al. Ultraconservation identifies a small subset of extremely constrained developmental enhancers. Nature Genetics, 2008, 40(2): 158–160.

85. Wang Y Zhao Y, Wang Y, Li Z, Guo B and Merilä J. 2020, Population transcriptomics reveals weak parallel genetic basis in repeated marine and freshwater divergence in nine-spined sticklebacks. Molecular Ecology. Accepted Author Manuscript. doi:10.1111/mec.15435

86. Wassarman PM, Jovine L, Litscher ES (2001) A profile of fertilization in mammals. Nature Cell Biology, 2001, 3(2): E59–E64.

87. Zhao H, Sun Z, Wang J, et al. CrossMap: a versatile tool for coordinate conversion between genome assemblies. Bioinformatics, 2014, 30(7): 1006–1007.

88. Zhu CD, Wang ZH, Yan B. Strategies for hypoxia adaptation in fish species: a review. Journal of Comparative Physiology B, 2013, 183(8): 1005–1013.

