## Supplementary Information for "We shall meet again - Genomics of historical admixture in the sea"

### Content

#### Supplementary Figures

Supplementary Figure S1. The first two principal components of the PCA analysis with all individuals from each nine-spined stickleback population.

Supplementary Figure S2. The 25 introgression-enriched regions identified with nine pairwise *fd* analyses.

Supplementary Figure S3. Correlations between mean *fd* and estimates of population differentiation ($F_{ST}$), absolute divergence ($d_{xy}$), and nucleotide diversity ($\pi$) of the three northern Baltic Sea populations.

#### Supplementary Tables

Supplementary Table S1. Detailed sample information.

Supplementary Table S2. Details of data filtering for each analysis

Supplementary Table S3. D-statistic of introgression tests among populations.

Supplementary Table S4. Results of f4-ratio test of ancestry proportion among different sets of populations.

Supplementary Table S5. Result of f4-ratio test and enrichment or depletion of ancestry proportion (alpha) in different genomic categories of admixed populations.

Supplementary Table S6. Test of ILS and detailed information for each candidate region identified by fd analysis.

Supplementary Table S7. Candidate regions identified by U and Q95 test, and genes in each region.

Supplementary Table S8. Classification of SNPs in NBS and SWE-BYN population.

Supplementary Table S9. Spearman rank correlations (rs) between admixture proportion (fd) and recombination rate for each admixed population.

Supplementary Table S10. Placement of the randomly selected representative individuals within the phylogenetic trees.

### Supplementary Figures

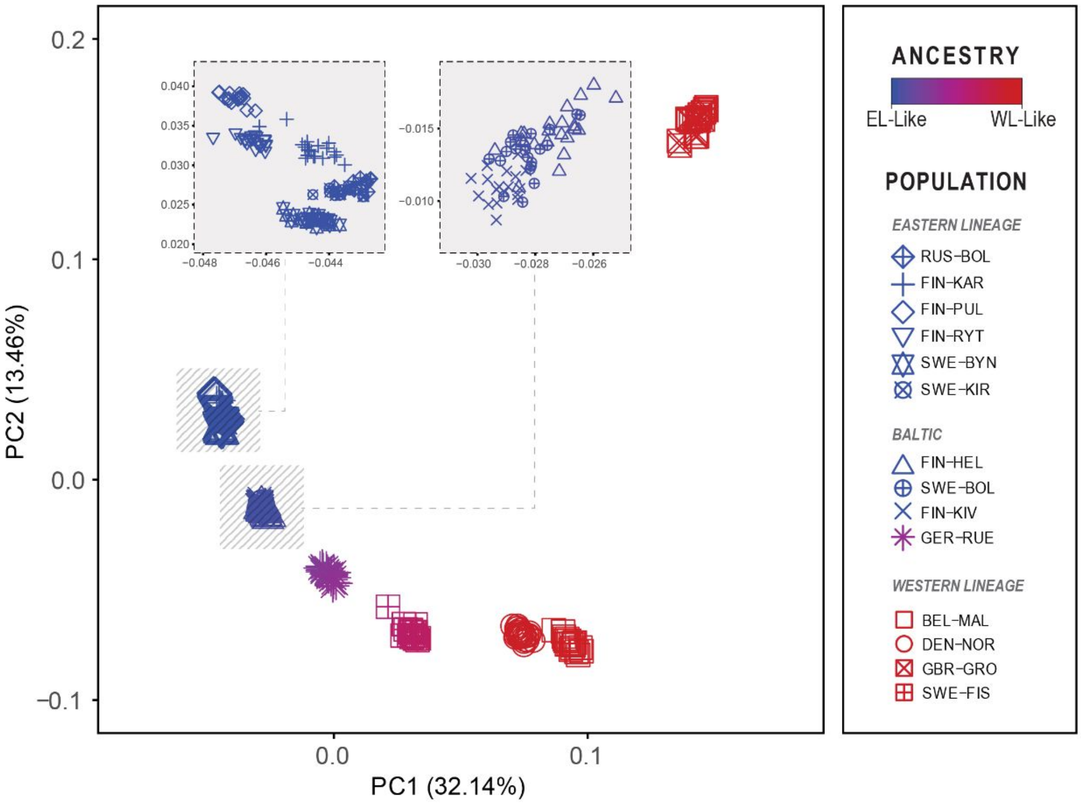


**Supplementary Figure S1.** The first two principal components of the PCA analysis with all individuals from each nine-spined stickleback population. The colour indicates the proportion of “EL-like” ancestry as estimated with ADMIXTURE (K=2).

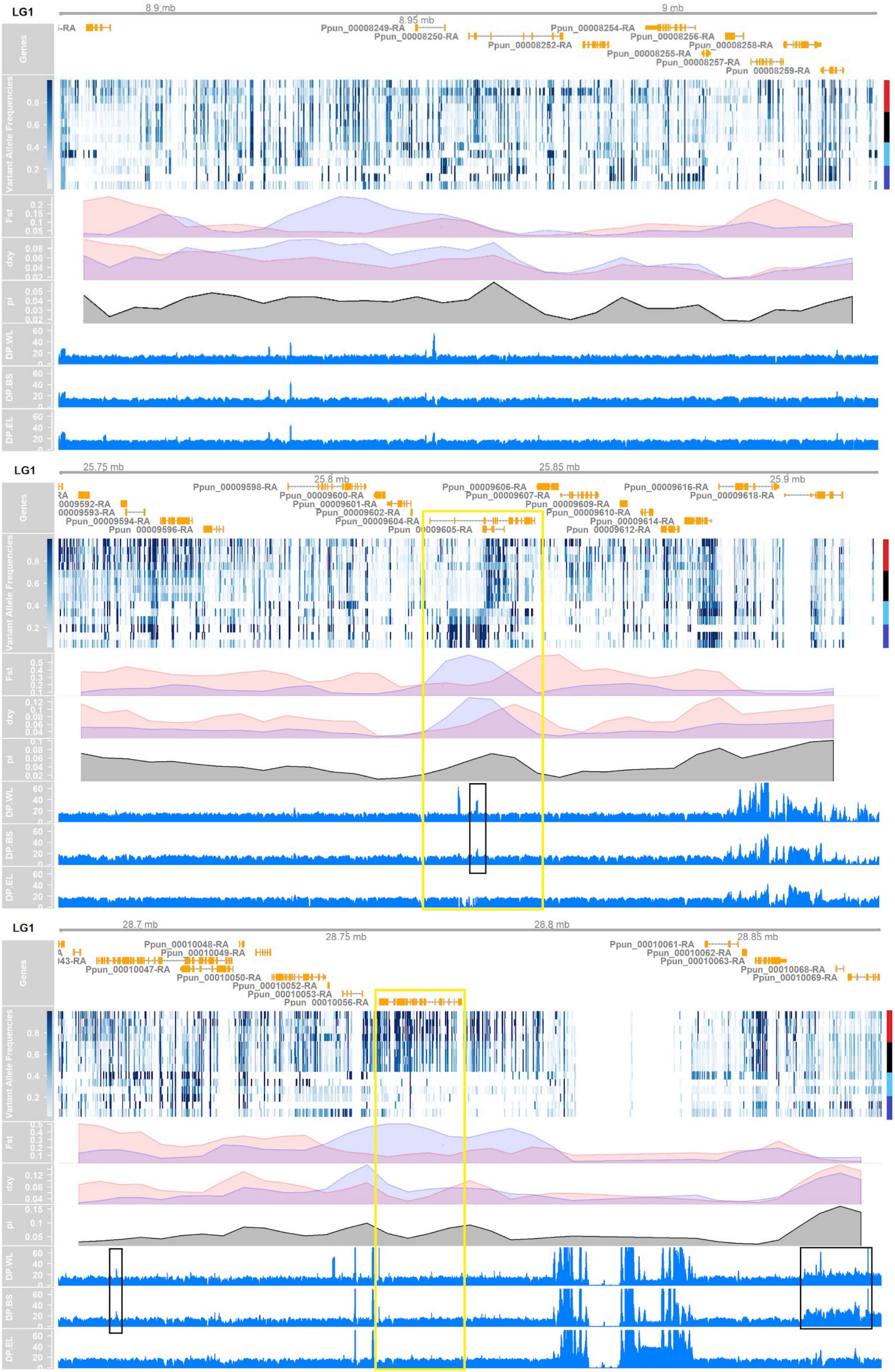

LG1
8.9 mb
8.95 mb
9 mb
Ppun_00008249-RA
Ppun_00008250-RA
Ppun_00008252-RA
Ppun_00008254-RA
Ppun_00008255-RA
Ppun_00008256-RA
Ppun_00008257-RA
Ppun_00008258-RA
Genes
Variant Allele Frequencies
Fst
dxy
pi
DP.WL
DP.BS
DP.EL
LG1
25.75 mb
25.8 mb
25.85 mb
25.9 mb
Ppun_00009598-RA
Ppun_00009600-RA
Ppun_00009601-RA
Ppun_00009602-RA
Ppun_00009604-RA
Ppun_00009606-RA
Ppun_00009607-RA
Ppun_00009609-RA
Ppun_00009610-RA
Ppun_00009614-RA
Ppun_00009616-RA
Ppun_00009618-RA
Ppun_00009594-RA
Ppun_00009596-RA
LG1
28.7 mb
28.75 mb
28.8 mb
28.85 mb
Ppun_00010047-RA
Ppun_00010048-RA
Ppun_00010049-RA
Ppun_00010050-RA
Ppun_00010052-RA
Ppun_00010053-RA
Ppun_00010056-RA
Ppun_00010061-RA
Ppun_00010062-RA
Ppun_00010063-RA
Ppun_00010068-RA
Ppun_00010069-RA

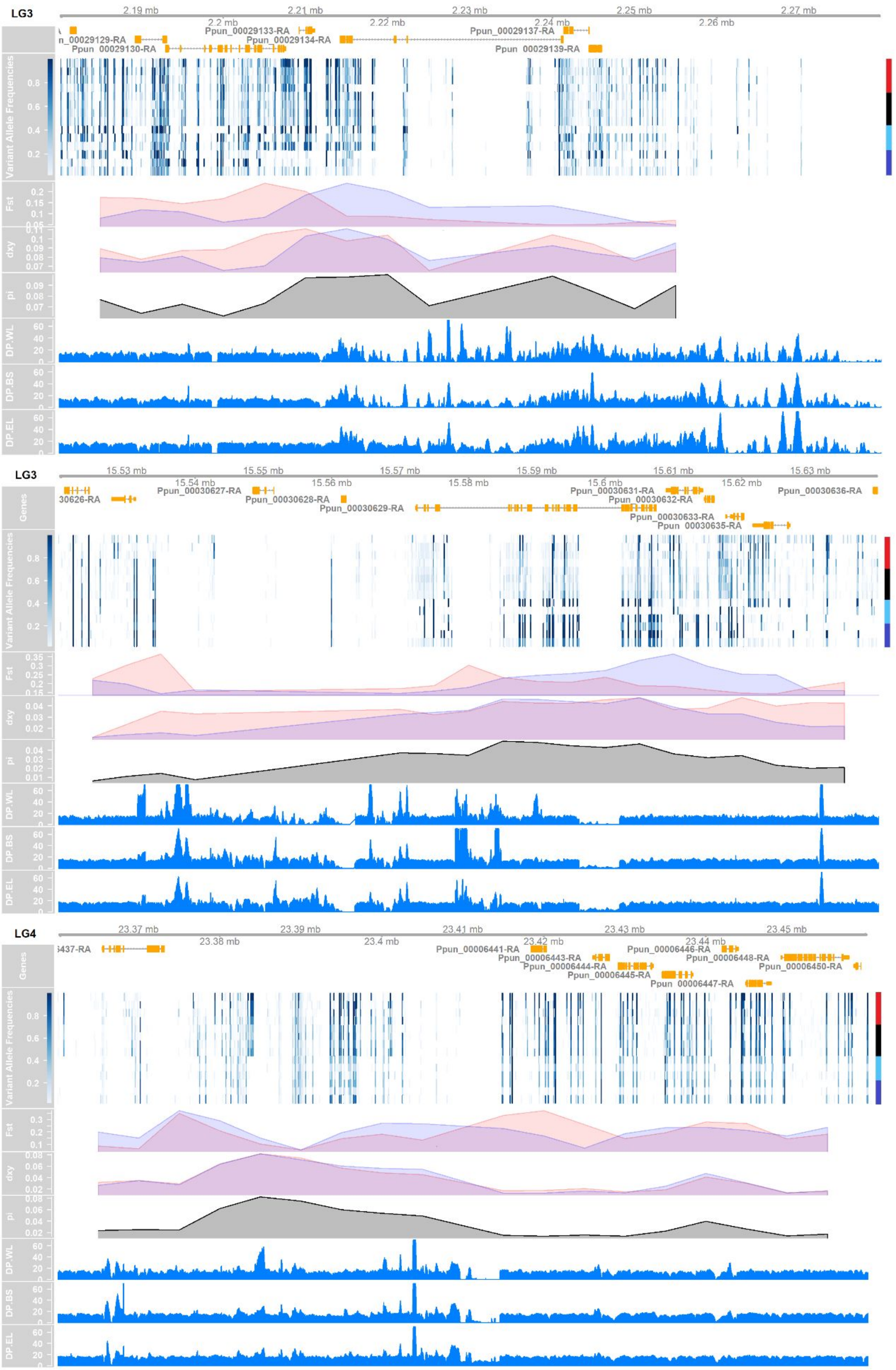
LG3
2.19 mb
2.2 mb
2.21 mb
2.22 mb
2.23 mb
2.24 mb
2.25 mb
2.26 mb
2.27 mb
Ppun_00029133-RA
Ppun_00029137-RA
n_00029129-RA
Ppun_00029134-RA
Ppun_00029130-RA
Ppun_00029139-RA
Variant Allele Frequencies
Fst
dxy
pi
DP.WL
DP.BS
DP.EL
LG3
15.53 mb
15.54 mb
15.55 mb
15.56 mb
15.57 mb
15.58 mb
15.59 mb
15.6 mb
15.61 mb
15.62 mb
15.63 mb
Genes
Ppun_00030627-RA
Ppun_00030631-RA
Ppun_00030636-RA
30626-RA
Ppun_00030628-RA
Ppun_00030632-RA
Ppun_00030629-RA
Ppun_00030633-RA
Ppun_00030635-RA
Variant Allele Frequencies
Fst
dxy
pi
DP.WL
DP.BS
DP.EL
LG4
23.37 mb
23.38 mb
23.39 mb
23.4 mb
23.41 mb
23.42 mb
23.43 mb
23.44 mb
23.45 mb
Genes
i437-RA
Ppun_00006441-RA
Ppun_00006446-RA
Ppun_00006443-RA
Ppun_00006448-RA
Ppun_00006444-RA
Ppun_00006450-RA
Ppun_00006445-RA
Ppun_00006447-RA
Variant Allele Frequencies
Fst
dxy
pi
DP.WL
DP.BS
DP.EL

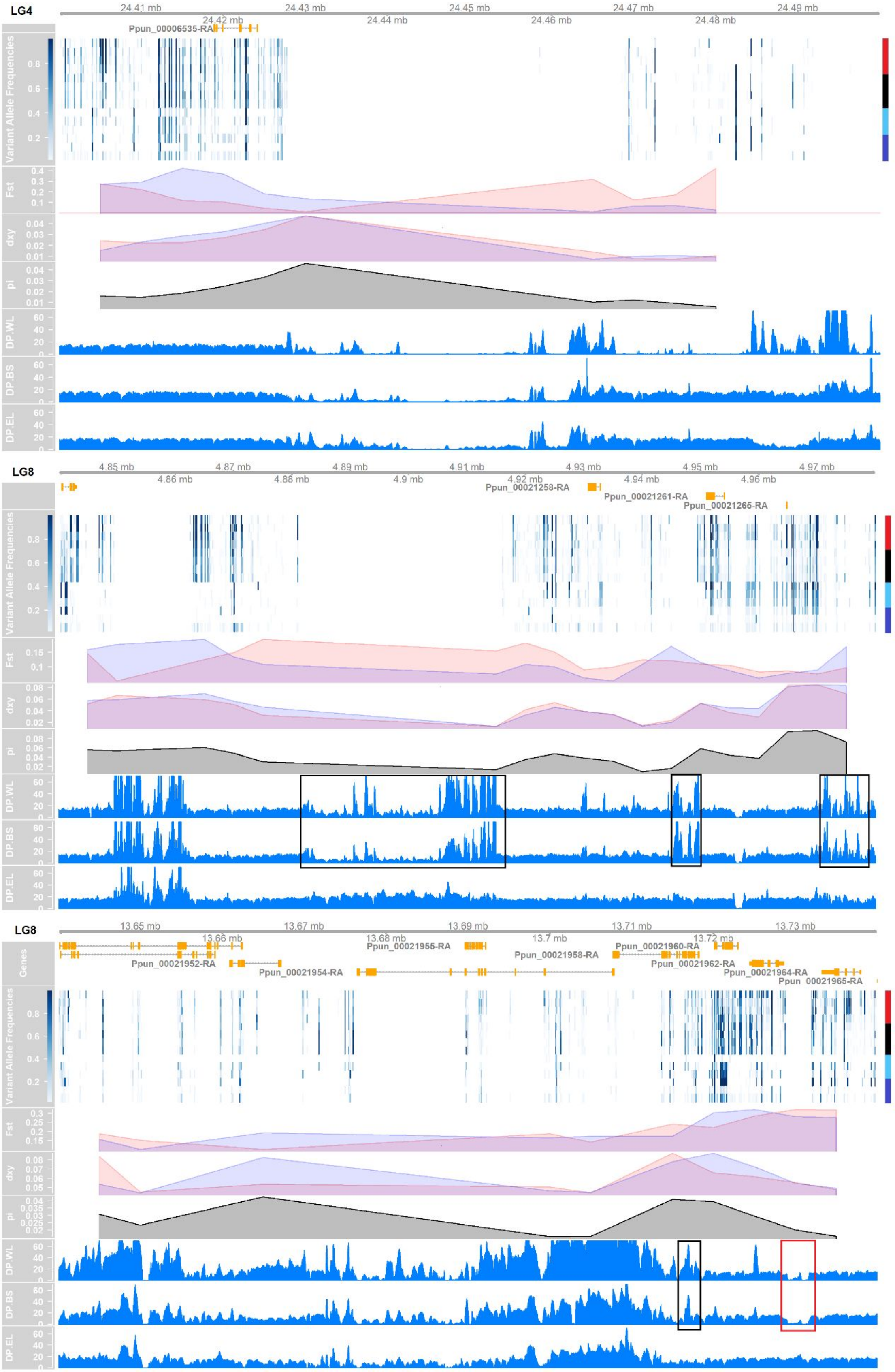

LG4
24.41 mb
24.42 mb
24.43 mb
24.44 mb
24.45 mb
24.46 mb
24.47 mb
24.48 mb
24.49 mb
Ppun_00006535-RA
Variant Allele Frequencies
Fst
dxy
pi
DP.WL
DP.BS
DP.EL
LG8
4.85 mb
4.86 mb
4.87 mb
4.88 mb
4.89 mb
4.9 mb
4.91 mb
4.92 mb
4.93 mb
4.94 mb
4.95 mb
4.96 mb
4.97 mb
Ppun_00021258-RA
Ppun_00021261-RA
Ppun_00021265-RA
LG8
13.65 mb
13.66 mb
13.67 mb
13.68 mb
13.69 mb
13.7 mb
13.71 mb
13.72 mb
13.73 mb
Genes
Ppun_00021952-RA
Ppun_00021954-RA
Ppun_00021955-RA
Ppun_00021958-RA
Ppun_00021960-RA
Ppun_00021962-RA
Ppun_00021964-RA
Ppun_00021965-RA

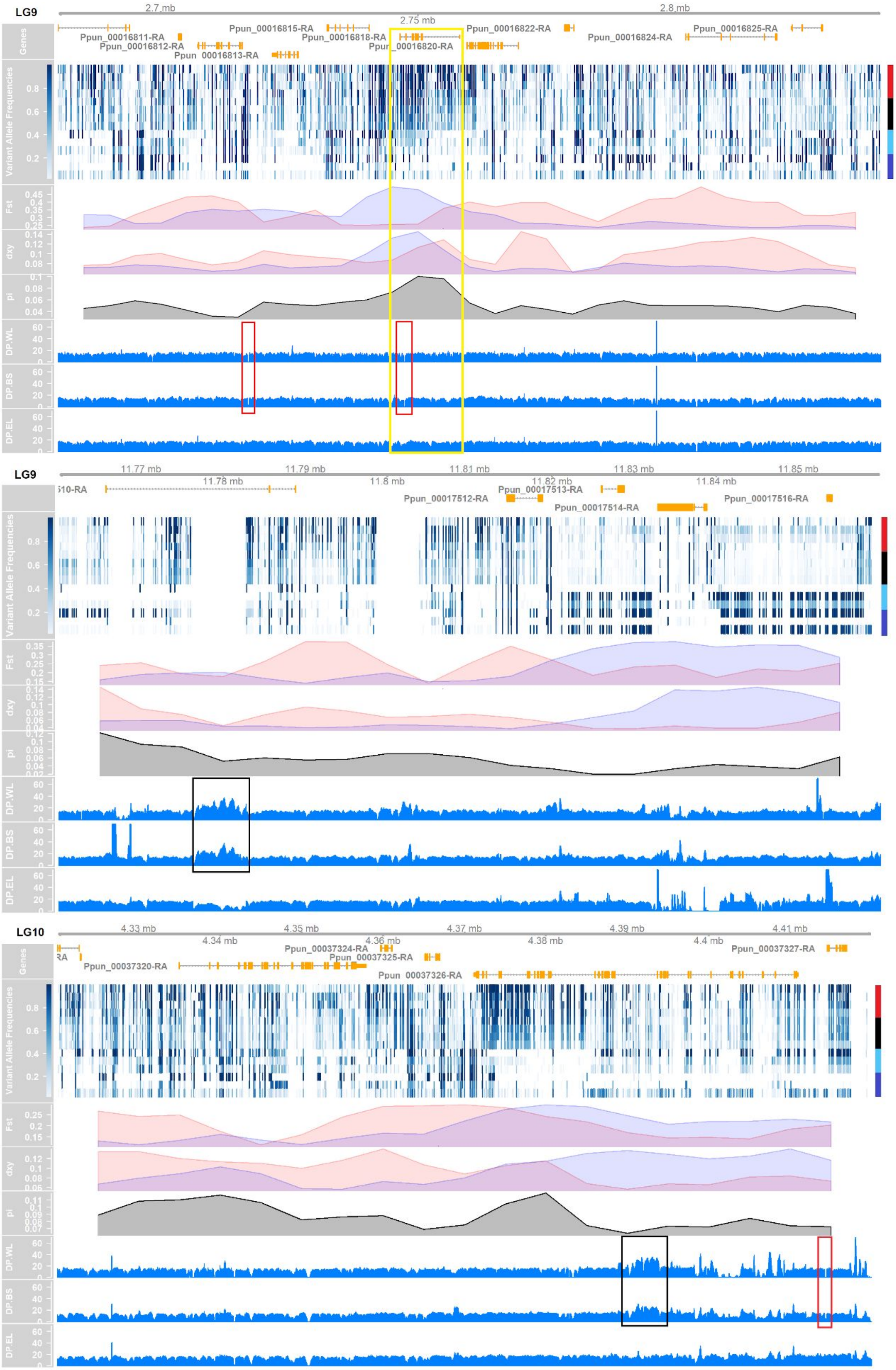

LG9
2.7 mb
2.75 mb
2.8 mb
Genes
Ppun_00016811-RA
Ppun_00016812-RA
Ppun_00016813-RA
Ppun_00016815-RA
Ppun_00016818-RA
Ppun_00016820-RA
Ppun_00016822-RA
Ppun_00016824-RA
Ppun_00016825-RA
Variant Allele Frequencies
Fst
dxy
pi
DP.WL
DP.BS
DP.EL
LG9
11.77 mb
11.78 mb
11.79 mb
11.8 mb
11.81 mb
11.82 mb
11.83 mb
11.84 mb
11.85 mb
Ppun_00017512-RA
Ppun_00017513-RA
Ppun_00017514-RA
Ppun_00017516-RA
Variant Allele Frequencies
Fst
dxy
pi
DP.WL
DP.BS
DP.EL
LG10
4.33 mb
4.34 mb
4.35 mb
4.36 mb
4.37 mb
4.38 mb
4.39 mb
4.4 mb
4.41 mb
Genes
Ppun_00037320-RA
Ppun_00037324-RA
Ppun_00037325-RA
Ppun_00037326-RA
Ppun_00037327-RA
Variant Allele Frequencies
Fst
dxy
pi
DP.WL
DP.BS
DP.EL

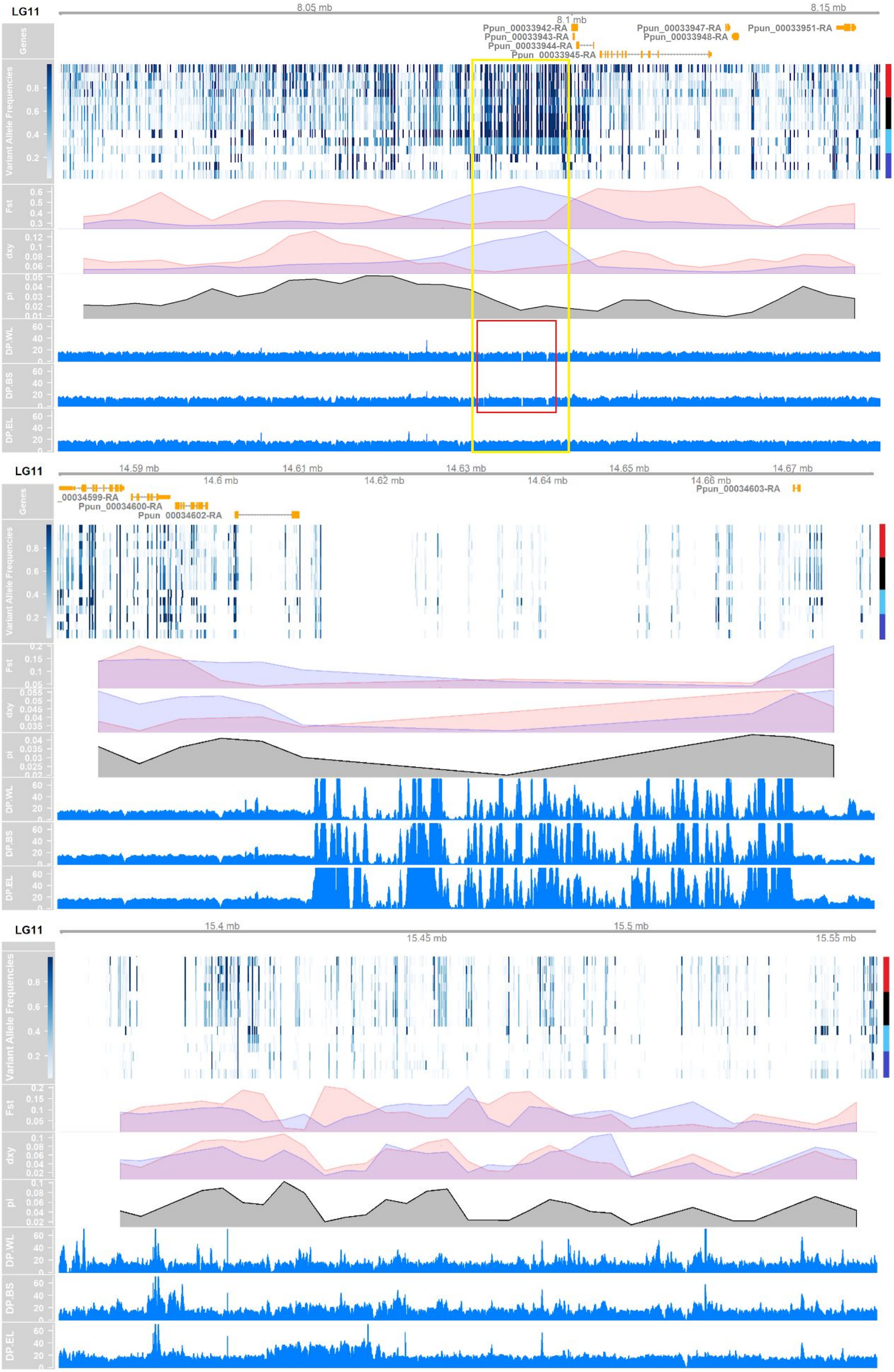

LG11
8.05 mb
8.1 mb
8.15 mb
Genes
Ppun_00033942-RA
Ppun_00033943-RA
Ppun_00033944-RA
Ppun_00033945-RA
Ppun_00033947-RA
Ppun_00033948-RA
Ppun_00033951-RA
Variant Allele Frequencies
Fst
dxy
pi
DP.WL
DP.BS
DP.EL
LG11
14.59 mb
14.6 mb
14.61 mb
14.62 mb
14.63 mb
14.64 mb
14.65 mb
14.66 mb
14.67 mb
_00034599-RA
Ppun_00034600-RA
Ppun_00034602-RA
Ppun_00034603-RA
LG11
15.4 mb
15.45 mb
15.5 mb
15.55 mb

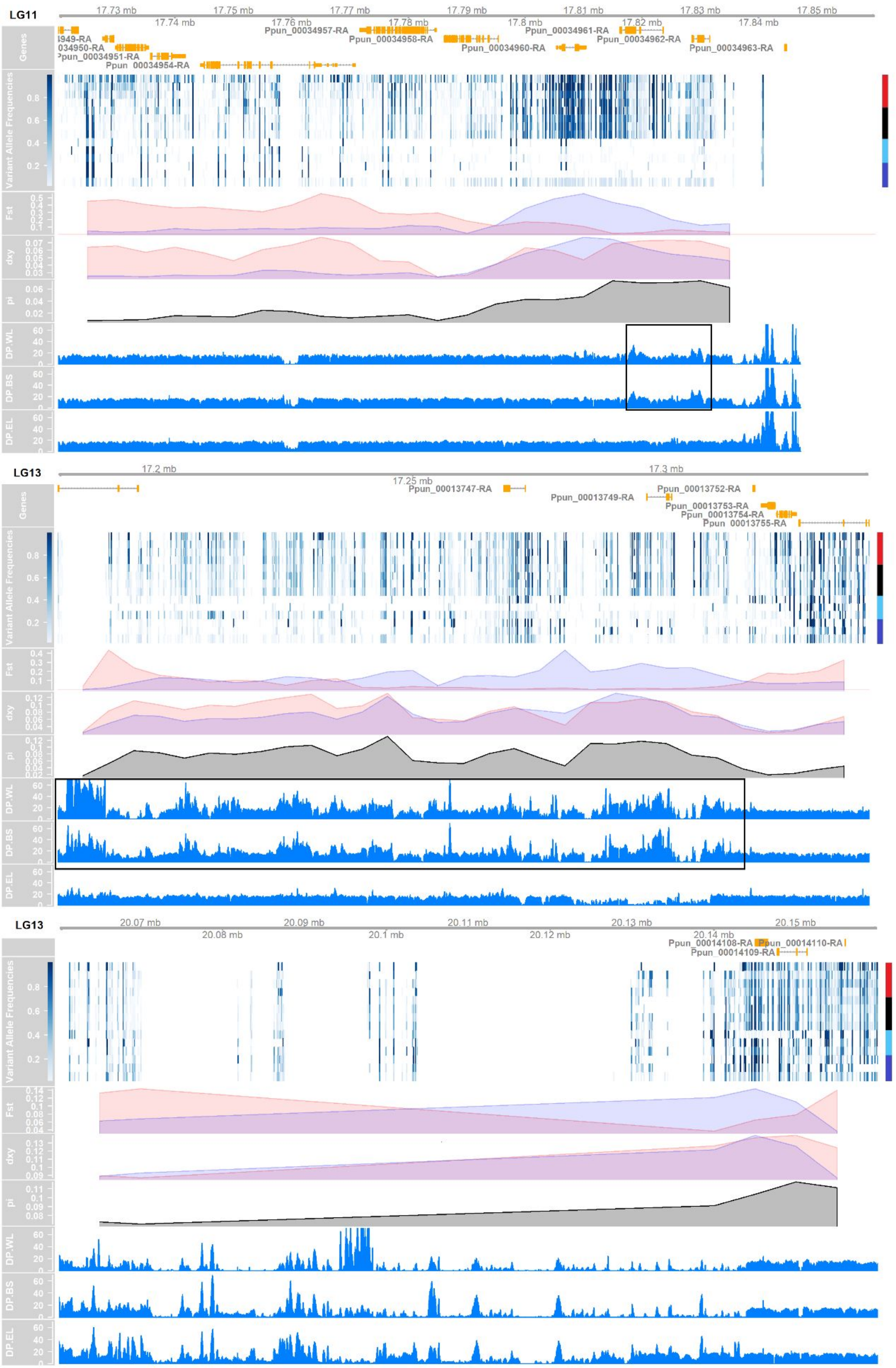

LG11
17.73 mb
17.74 mb
17.75 mb
17.76 mb
17.77 mb
17.78 mb
17.79 mb
17.8 mb
17.81 mb
17.82 mb
17.83 mb
17.84 mb
17.85 mb
Genes
Ppun_00034957-RA
Ppun_00034958-RA
Ppun_00034960-RA
Ppun_00034961-RA
Ppun_00034962-RA
Ppun_00034963-RA
034950-RA
Ppun_00034954-RA
Variant Allele Frequencies
Fst
dxy
pi
DP.WL
DP.BS
DP.EL
LG13
17.2 mb
17.25 mb
17.3 mb
Ppun_00013747-RA
Ppun_00013749-RA
Ppun_00013752-RA
Ppun_00013753-RA
Ppun_00013754-RA
Ppun_00013755-RA
LG13
20.07 mb
20.08 mb
20.09 mb
20.1 mb
20.11 mb
20.12 mb
20.13 mb
20.14 mb
20.15 mb
Ppun_00014108-RA
Ppun_00014109-RA
Ppun_00014110-RA

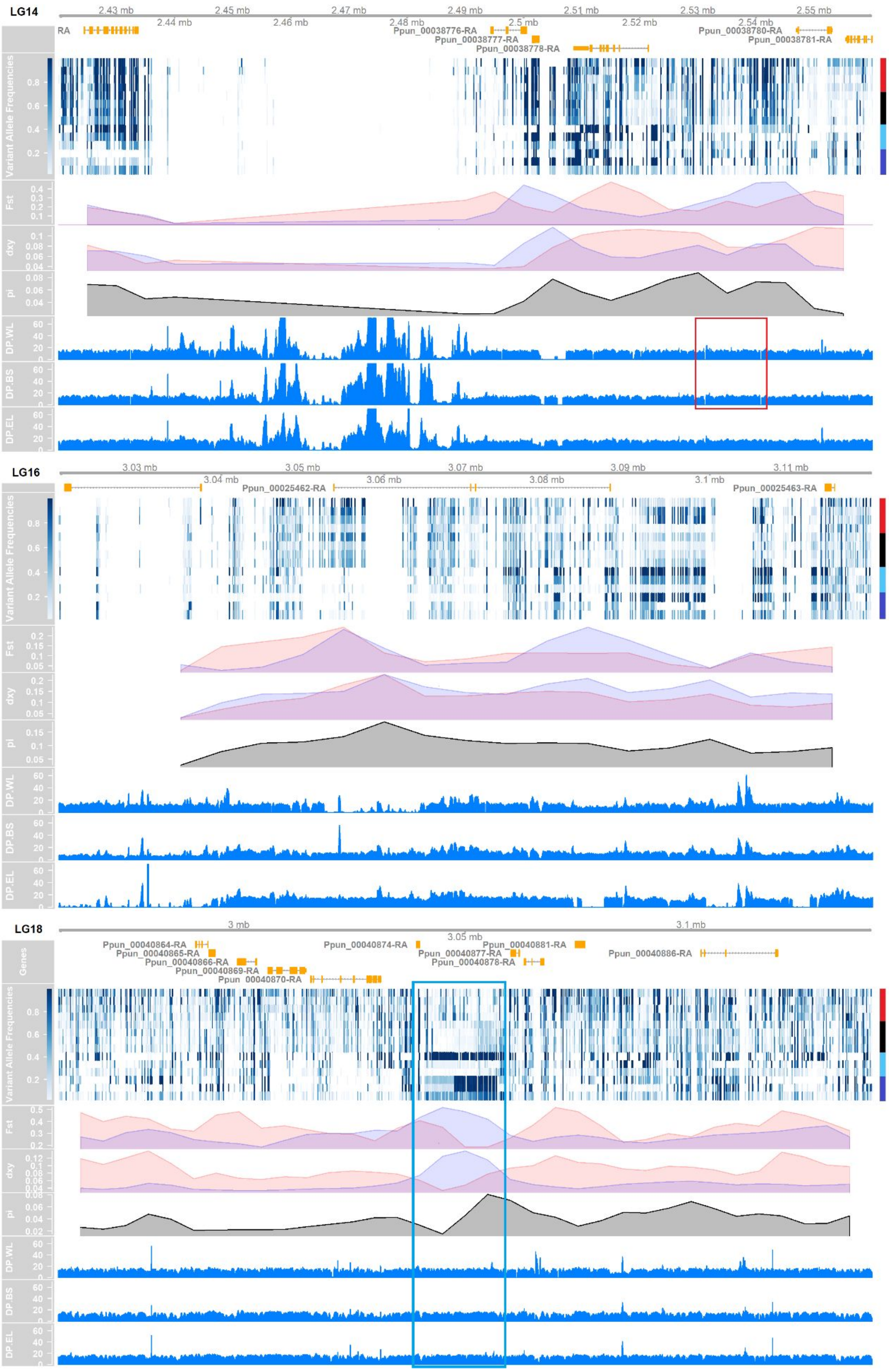
LG14
2.43 mb
2.44 mb
2.45 mb
2.46 mb
2.47 mb
2.48 mb
2.49 mb
2.5 mb
2.51 mb
2.52 mb
2.53 mb
2.54 mb
2.55 mb
RA
Ppun_00038776-RA
Ppun_00038777-RA
Ppun_00038778-RA
Ppun_00038780-RA
Ppun_00038781-RA
Variant Allele Frequencies
Fst
dxy
pi
DP.WL
DP.BS
DP.EL
LG16
3.03 mb
3.04 mb
3.05 mb
3.06 mb
3.07 mb
3.08 mb
3.09 mb
3.1 mb
3.11 mb
Ppun_00025462-RA
Ppun_00025463-RA
LG18
3 mb
3.05 mb
3.1 mb
Genes
Ppun_00040864-RA
Ppun_00040865-RA
Ppun_00040866-RA
Ppun_00040869-RA
Ppun_00040870-RA
Ppun_00040874-RA
Ppun_00040877-RA
Ppun_00040878-RA
Ppun_00040881-RA
Ppun_00040886-RA

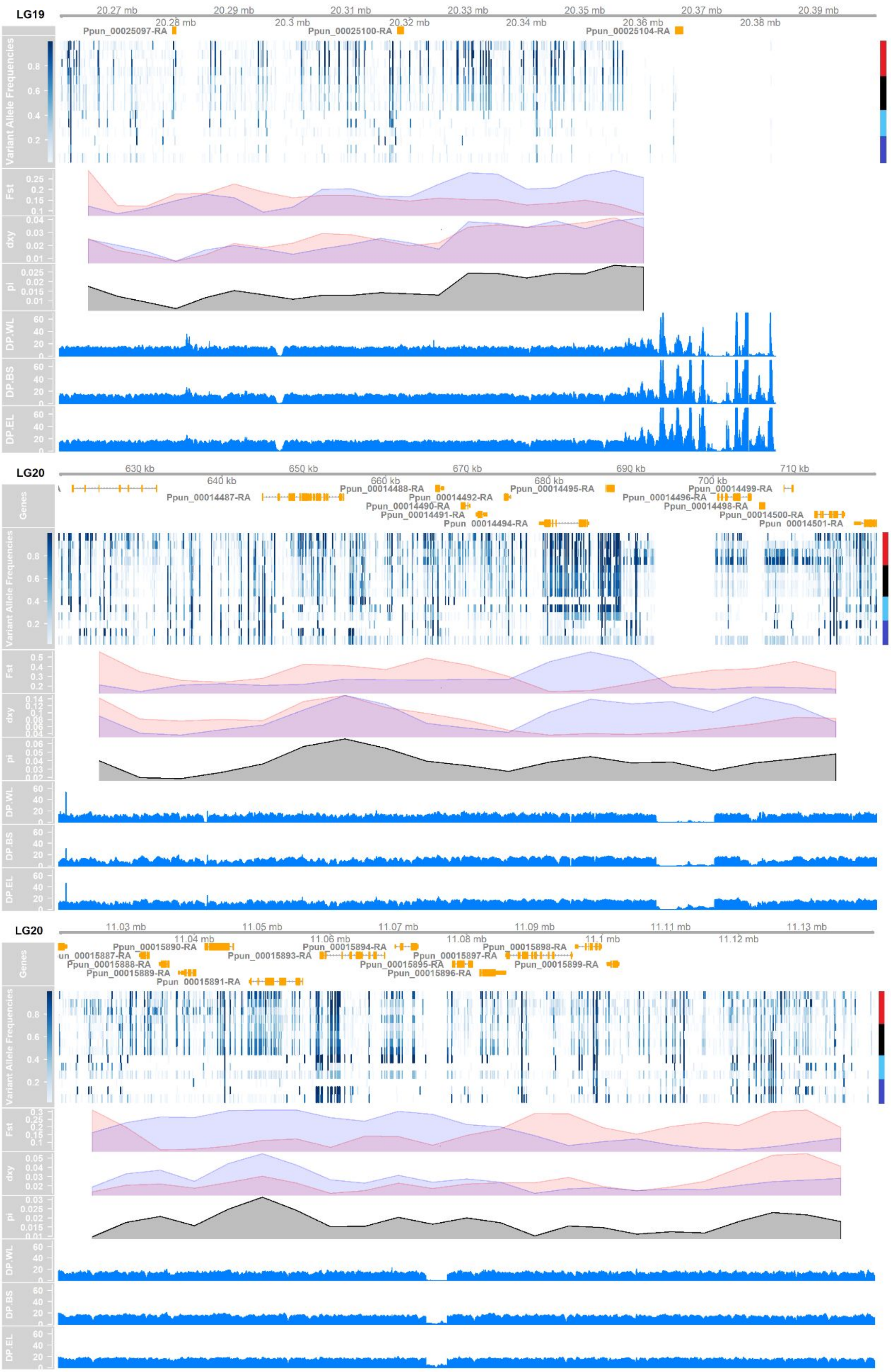

LG19
20.27 mb
20.28 mb
20.29 mb
20.3 mb
20.31 mb
20.32 mb
20.33 mb
20.34 mb
20.35 mb
20.36 mb
20.37 mb
20.38 mb
20.39 mb
Ppun_00025097-RA
Ppun_00025100-RA
Ppun_00025104-RA
Variant Allele Frequencies
Fst
dxy
pi
DP.WL
DP.BS
DP.EL
LG20
630 kb
640 kb
650 kb
660 kb
670 kb
680 kb
690 kb
700 kb
710 kb
Genes
Ppun_00014487-RA
Ppun_00014488-RA
Ppun_00014490-RA
Ppun_00014491-RA
Ppun_00014492-RA
Ppun_00014494-RA
Ppun_00014495-RA
Ppun_00014496-RA
Ppun_00014498-RA
Ppun_00014499-RA
Ppun_00014500-RA
Ppun_00014501-RA
LG20
11.03 mb
11.04 mb
11.05 mb
11.06 mb
11.07 mb
11.08 mb
11.09 mb
11.1 mb
11.11 mb
11.12 mb
11.13 mb
Ppun_00015887-RA
Ppun_00015888-RA
Ppun_00015889-RA
Ppun_00015890-RA
Ppun_00015891-RA
Ppun_00015893-RA
Ppun_00015894-RA
Ppun_00015895-RA
Ppun_00015896-RA
Ppun_00015897-RA
Ppun_00015898-RA
Ppun_00015899-RA

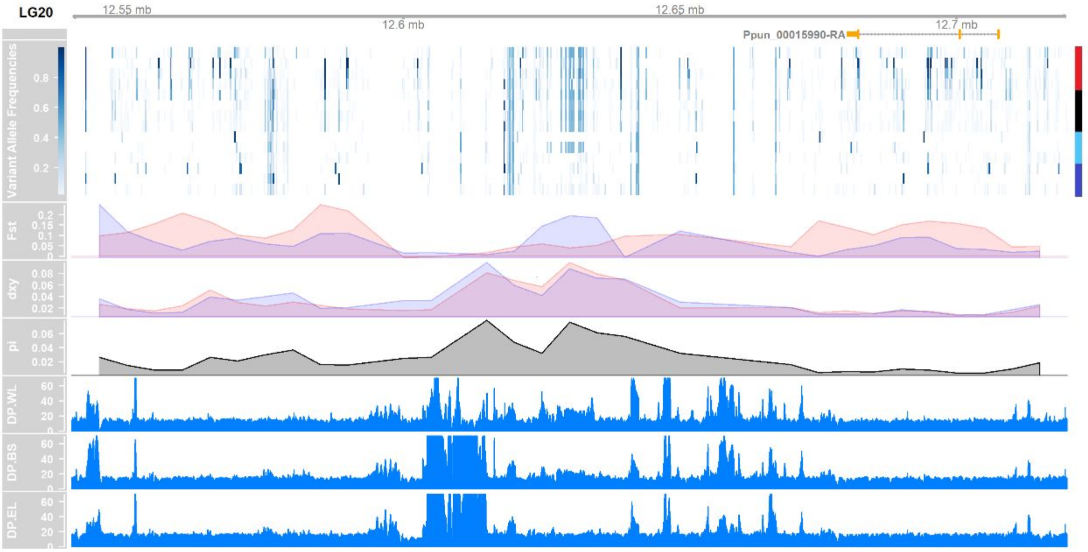


**Supplementary Figure S2.** 25 introgression-enriched regions identified with nine pairwise *fd* analyses (Methods). The tracks show gene structure (coding sequences in orange), per site variant allele frequencies (heatmap), $F_{ST}$, $d_{xy}$, and $\pi$ variation in the three northern Baltic Sea populations (NBS; i.e. FIN-HEL, SWE-BOL, and FIN-KIV), and mean sequencing depth in WL (GBR-GRO), NBS, and EL (RUS-BOL). Color gradient indicates the variant allele frequency from low (white) to high (dark blue). Red, black, light blue, and blue bars on the right refer to WL, Baltic Sea, EL (admixed), EL (non-admixed) populations, respectively, in the order GBR-GRO, BEL-MAL, DEN-NOR, SWE-FIS, GER-RUE, FIN-HEL, SWE-BOL, FIN-KIV, SWE-BYN, SWE-KIR, FIN-KAR, FIN-PUL, FIN-RYT, and RUS-BOL. $F_{ST}$ and $d_{xy}$ computed against RUS-BOL and GBR-GRO are shown in light blue and pink, respectively. Black and red squares highlight genomic regions containing duplicates and deletions uniquely shared between WL and NBS. Light blue square highlights the putative inversion shared by the three EL (non-admixed) populations in LG18. Yellow squares highlight the inferred adaptively introgressed regions.

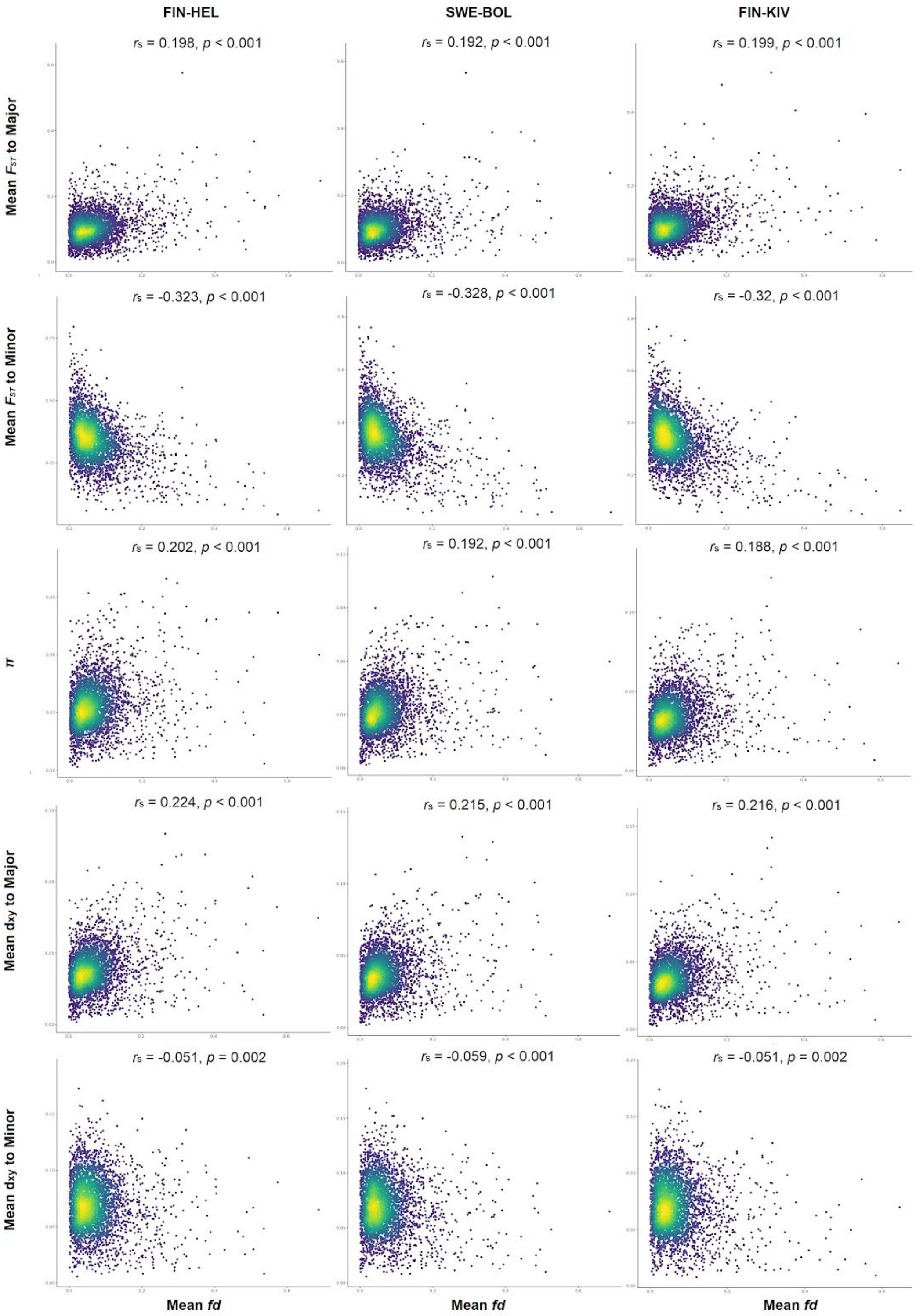


**Supplementary Figure S3.** Correlations between mean *fd* (mean of nine pair-wise estimates) and estimates of population differentiation ($F_{ST}$), absolute divergence ($d_{xy}$), and nucleotide diversity ($\pi$) of the three northern Baltic Sea populations (FIN-HEL, SWE-BOL, FIN-KIV).

Table S1. Information of the Pungitius pungitius and outgroup samples used. N = number of individuals used.

| Abbreviation | Species | Country | Locality | Latitude | Longitude | Habitat | N | Lineage | Year of Sampling |
|---|---|---|---|---|---|---|---|---|---|
| BEL-MAL | *Pungitius pungitius* | Belgium | Maldegem | 51.17473056 | 3.469569444 | Freshwater | 20 | European Western Lineage | 2011 |
| DEN-NOR | *Pungitius pungitius* | Denmark | Emmerlev Klev | 54.98482778 | 8.661875 | Marine | 25 | European Western Lineage | 2011 |
| SWE-FIS | *Pungitius pungitius* | Sweden | Fiskebäckskil, Väster Götaland | 58.23333333 | 11.4 | Marine | 20 | European Western Lineage | 2009 |
| GBR-GRO | *Pungitius pungitius* | United Kindom | Loch Grogarry, Scotland | 57.61512222 | -7.511461111 | Freshwater | 19 | European Western Lineage | 2015 |
| FIN-HEL | *Pungitius pungitius* | Finland | Uutela, Helsinki | 60.2025 | 25.18277778 | Marine | 22 | European Eastern Lineage | 2013 |
| FIN-RYT | *Pungitius pungitius* | Finland | Rytilampi, Kuusamo | 66.38416667 | 29.32 | Freshwater | 21 | European Eastern Lineage | 2006 |
| FIN-KIV | *Pungitius pungitius* | Finland | Kiviniemi | 65 | 25.46666667 | Marine | 19 | European Eastern Lineage | 2003 |
| FIN-PUL | *Pungitius pungitius* | Finland | Pulmankijärvi, Kevo | 69.96666667 | 27.96666667 | Freshwater | 17 | European Eastern Lineage | 2006 |
| FIN-KAR | *Pungitius pungitius* | Finland | Karhulampi, Rovaniemi | 66.65666667 | 26.44083333 | Freshwater | 20 | European Eastern Lineage | 2009 |
| GER-RUE | *Pungitius pungitius* | Germany | Rügen | 54.00667778 | 13.00353333 | Marine | 27 | European Eastern Lineage | 2009 |
| RUS-BOL | *Pungitius pungitius* | Russia | Bolotnoje, Chkalov | 66.3 | 33.4 | Freshwater | 20 | European Eastern Lineage | 2010 |
| SWE-BOL | *Pungitius pungitius* | Sweden | Bölesviken | 63.66138889 | 20.21194444 | Marine | 21 | European Eastern Lineage | 2007 |
| SWE-BYN | *Pungitius pungitius* | Sweden | Bynästjärnen, Amsele | 64.45527778 | 19.44444444 | Freshwater | 23 | European Eastern Lineage | 2013 |
| SWE-KIR | *Pungitius pungitius* | Sweden | Kiruna | 67.89611111 | 20.08611111 | Freshwater | 15 | European Eastern Lineage | 2002 |
| PUN-TYM | *Pungitius tymensis* | Japan | Motosakimui River, Hokkaido | 43.82777778 | 145.0861111 | Freshwater | 1 | Outgroup species | 2007 |

| Table S2. Details of data filtering for each analysis | | |
|---|---|---|
| **Name** | **Filtering Criteria** | **Used in analysis** |
| SNP Set 0 | Original dataset, removing identified repetitive sequences and interspecific variants, only autosomal biallelic SNPs retained | SMC++ |
| SNP Set 1 | SNP Set 0, keep only binary SNPs with quanlity score ≥ 30, mean coverage ≥ 5x and ≤ 25x. | *NA* |
| SNP Set 2 | SNP Set 1, allowing maximum 25% missing data, mean coverage ≥ 6.5x and ≤ 20x, distance between two SNPs at least 10kb. | PCA, ADMIXTURE |
| SNP Set 3 | SNP Set 1, allowing maximum 10% missing data. | ADMIXTOOLS (D statistic and f4-ratio test) |
| SNP Set 4 | SNP Set 1, allowing maximum 20% missing data. | *fd**, summary statistics**, per-site FST estimation |
| SNP Set 5 | SNP Set 1, allowing maximum 25% missing data. | U test, Q95 test and SNP classification *** |

* In fd calculation, missing data was controlled with the ABBABABAwindows.py script by setting the parameter --minData 0.8.

** Missing data was controlled by using vcftools --max-missing 0.8 then use popgenWindows.py script to calculate summary statistics Fst dxy and pi.

*** Controlling missing data in U test, Q95 test and SNP classificaiton was done with costum R script.

Table S3. D-statistic of introgression tests among populations.

| P1 | P2 | P3 | O | D | std(D) | Z | nBABA | nABBA |
|---|---|---|---|---|---|---|---|---|
| DEN-NOR | GBR-GRO | RUS-BOL | PUN-TYM | 0.2683 | 0.004384 | 61.207 | 111518 | 64333 |
| DEN-NOR | GBR-GRO | FIN-RYT | PUN-TYM | 0.2692 | 0.004378 | 61.494 | 111476 | 64185 |
| DEN-NOR | GBR-GRO | FIN-PUL | PUN-TYM | 0.2665 | 0.0045 | 59.235 | 111335 | 64474 |
| SWE-FIS | GBR-GRO | RUS-BOL | PUN-TYM | 0.3943 | 0.00444 | 88.809 | 145535 | 63228 |
| SWE-FIS | GBR-GRO | FIN-RYT | PUN-TYM | 0.3933 | 0.004412 | 89.141 | 145206 | 63237 |
| SWE-FIS | GBR-GRO | FIN-PUL | PUN-TYM | 0.3921 | 0.004633 | 84.633 | 145233 | 63423 |
| BEL-MAL | GBR-GRO | RUS-BOL | PUN-TYM | 0.2182 | 0.004775 | 45.689 | 100613 | 64576 |
| BEL-MAL | GBR-GRO | FIN-RYT | PUN-TYM | 0.2198 | 0.004767 | 46.102 | 100632 | 64370 |
| BEL-MAL | GBR-GRO | FIN-PUL | PUN-TYM | 0.2181 | 0.004863 | 44.855 | 100619 | 64584 |
| GBR-GRO | BEL-MAL | RUS-BOL | PUN-TYM | -0.2182 | 0.004775 | -45.689 | 64576 | 100613 |
| GBR-GRO | BEL-MAL | FIN-RYT | PUN-TYM | -0.2198 | 0.004767 | -46.102 | 64370 | 100632 |
| GBR-GRO | BEL-MAL | FIN-PUL | PUN-TYM | -0.2181 | 0.004863 | -44.855 | 64584 | 100619 |
| GER-RUE | RUS-BOL | GBR-GRO | PUN-TYM | 0.2665 | 0.003909 | 68.159 | 103568 | 59990 |
| FIN-HEL | RUS-BOL | GBR-GRO | PUN-TYM | 0.1291 | 0.003735 | 34.559 | 71813 | 55396 |
| SWE-BOL | RUS-BOL | GBR-GRO | PUN-TYM | 0.1246 | 0.003486 | 35.744 | 70633 | 54982 |
| FIN-KIV | RUS-BOL | GBR-GRO | PUN-TYM | 0.1181 | 0.003543 | 33.332 | 69802 | 55057 |
| SWE-BYN | RUS-BOL | GBR-GRO | PUN-TYM | 0.037 | 0.004428 | 8.347 | 53303 | 49504 |
| SWE-KIR | RUS-BOL | GBR-GRO | PUN-TYM | 0.0188 | 0.003942 | 4.764 | 50150 | 48303 |
| FIN-KAR | RUS-BOL | GBR-GRO | PUN-TYM | 0.0092 | 0.003055 | 3.001 | 47857 | 46988 |
| FIN-PUL | RUS-BOL | GBR-GRO | PUN-TYM | 0.0068 | 0.00351 | 1.937 | 46818 | 46186 |
| FIN-RYT | RUS-BOL | GBR-GRO | PUN-TYM | 0.006 | 0.003641 | 1.649 | 46555 | 46000 |
| GER-RUE | FIN-PUL | GBR-GRO | PUN-TYM | 0.2624 | 0.00429 | 61.164 | 103297 | 60359 |
| FIN-HEL | FIN-PUL | GBR-GRO | PUN-TYM | 0.1235 | 0.004164 | 29.651 | 71803 | 56023 |
| SWE-BOL | FIN-PUL | GBR-GRO | PUN-TYM | 0.1194 | 0.003936 | 30.329 | 70405 | 55391 |
| FIN-KIV | FIN-PUL | GBR-GRO | PUN-TYM | 0.1134 | 0.003857 | 29.396 | 69283 | 55176 |
| SWE-BYN | FIN-PUL | GBR-GRO | PUN-TYM | 0.0312 | 0.00489 | 6.374 | 52383 | 49218 |
| SWE-KIR | FIN-PUL | GBR-GRO | PUN-TYM | 0.0128 | 0.004254 | 3.001 | 48142 | 46930 |
| FIN-KAR | FIN-PUL | GBR-GRO | PUN-TYM | 0.0028 | 0.003885 | 0.708 | 43194 | 42958 |

| | | | | | | | | |
|---|---|---|---|---|---|---|---|---|
| FIN-RYT | FIN-PUL | GBR-GRO | PUN-TYM | -0.0008 | 0.004748 | -0.172 | 45457 | 45531 |
| RUS-BOL | FIN-PUL | GBR-GRO | PUN-TYM | -0.0068 | 0.00351 | -1.937 | 46186 | 46818 |
| GER-RUE | FIN-RYT | GBR-GRO | PUN-TYM | 0.2645 | 0.004174 | 63.372 | 102836 | 59815 |
| FIN-HEL | FIN-RYT | GBR-GRO | PUN-TYM | 0.1263 | 0.004177 | 30.234 | 70745 | 54883 |
| SWE-BOL | FIN-RYT | GBR-GRO | PUN-TYM | 0.1216 | 0.003924 | 31.004 | 69598 | 54503 |
| FIN-KIV | FIN-RYT | GBR-GRO | PUN-TYM | 0.1149 | 0.00404 | 28.448 | 68829 | 54639 |
| SWE-BYN | FIN-RYT | GBR-GRO | PUN-TYM | 0.0331 | 0.005041 | 6.564 | 50753 | 47502 |
| SWE-KIR | FIN-RYT | GBR-GRO | PUN-TYM | 0.0137 | 0.00498 | 2.744 | 47883 | 46592 |
| FIN-KAR | FIN-RYT | GBR-GRO | PUN-TYM | 0.0034 | 0.004054 | 0.843 | 46092 | 45778 |
| FIN-PUL | FIN-RYT | GBR-GRO | PUN-TYM | 0.0008 | 0.004748 | 0.172 | 45531 | 45457 |
| RUS-BOL | FIN-RYT | GBR-GRO | PUN-TYM | -0.006 | 0.003641 | -1.649 | 46000 | 46555 |

Table S4. Results of f4-ratio test of ancestry proportion among different sets of populations

| A | O | X | C | B | alpha | std(alpha) | Z | manner |
|---|---|---|---|---|---|---|---|---|
| FIN-PUL | PUN-TYM | DEN-NOR | GBR-GRO | RUS-BOL | 0.248912 | 0.003699 | 67.298 | Direct estimation of EL ancestries |
| FIN-PUL | PUN-TYM | SWE-FIS | GBR-GRO | RUS-BOL | 0.43455 | 0.004344 | 100.035 | Direct estimation of EL ancestries |
| FIN-PUL | PUN-TYM | BEL-MAL | GBR-GRO | RUS-BOL | 0.191404 | 0.003923 | 48.789 | Direct estimation of EL ancestries |
| FIN-PUL | PUN-TYM | GER-RUE | GBR-GRO | RUS-BOL | 0.607035 | 0.003639 | 166.833 | Direct estimation of EL ancestries |
| FIN-PUL | PUN-TYM | FIN-HEL | GBR-GRO | RUS-BOL | 0.774329 | 0.002934 | 263.959 | Direct estimation of EL ancestries |
| FIN-PUL | PUN-TYM | SWE-BOL | GBR-GRO | RUS-BOL | 0.785971 | 0.002753 | 285.502 | Direct estimation of EL ancestries |
| FIN-PUL | PUN-TYM | FIN-KIV | GBR-GRO | RUS-BOL | 0.798925 | 0.002772 | 288.228 | Direct estimation of EL ancestries |
| FIN-PUL | PUN-TYM | SWE-BYN | GBR-GRO | RUS-BOL | 0.94236 | 0.003801 | 247.915 | Direct estimation of EL ancestries |
| FIN-PUL | PUN-TYM | SWE-KIR | GBR-GRO | RUS-BOL | 0.991958 | 0.003182 | 311.789 | Direct estimation of EL ancestries |
| FIN-PUL | PUN-TYM | FIN-KAR | GBR-GRO | RUS-BOL | 1.068534 | 0.002867 | 372.724 | Direct estimation of EL ancestries |
| FIN-RYT | PUN-TYM | DEN-NOR | GBR-GRO | RUS-BOL | 0.25116 | 0.003662 | 68.592 | Direct estimation of EL ancestries |
| FIN-RYT | PUN-TYM | SWE-FIS | GBR-GRO | RUS-BOL | 0.435358 | 0.004187 | 103.984 | Direct estimation of EL ancestries |
| FIN-RYT | PUN-TYM | BEL-MAL | GBR-GRO | RUS-BOL | 0.193009 | 0.003904 | 49.435 | Direct estimation of EL ancestries |
| FIN-RYT | PUN-TYM | GER-RUE | GBR-GRO | RUS-BOL | 0.61506 | 0.003554 | 173.053 | Direct estimation of EL ancestries |
| FIN-RYT | PUN-TYM | FIN-HEL | GBR-GRO | RUS-BOL | 0.792349 | 0.003021 | 262.291 | Direct estimation of EL ancestries |
| FIN-RYT | PUN-TYM | SWE-BOL | GBR-GRO | RUS-BOL | 0.799393 | 0.002804 | 285.125 | Direct estimation of EL ancestries |
| FIN-RYT | PUN-TYM | FIN-KIV | GBR-GRO | RUS-BOL | 0.806515 | 0.002991 | 269.616 | Direct estimation of EL ancestries |
| FIN-RYT | PUN-TYM | SWE-BYN | GBR-GRO | RUS-BOL | 0.964335 | 0.004362 | 221.089 | Direct estimation of EL ancestries |
| FIN-RYT | PUN-TYM | SWE-KIR | GBR-GRO | RUS-BOL | 0.992527 | 0.003721 | 266.74 | Direct estimation of EL ancestries |
| FIN-RYT | PUN-TYM | FIN-KAR | GBR-GRO | RUS-BOL | 1.010425 | 0.0025 | 404.228 | Direct estimation of EL ancestries |
| FIN-PUL | PUN-TYM | DEN-NOR | GBR-GRO | FIN-RYT | 0.246783 | 0.003797 | 64.986 | Direct estimation of EL ancestries |
| FIN-PUL | PUN-TYM | SWE-FIS | GBR-GRO | FIN-RYT | 0.430837 | 0.004552 | 94.657 | Direct estimation of EL ancestries |
| FIN-PUL | PUN-TYM | BEL-MAL | GBR-GRO | FIN-RYT | 0.189758 | 0.003961 | 47.903 | Direct estimation of EL ancestries |
| FIN-PUL | PUN-TYM | GER-RUE | GBR-GRO | FIN-RYT | 0.601855 | 0.004097 | 146.884 | Direct estimation of EL ancestries |
| FIN-PUL | PUN-TYM | FIN-HEL | GBR-GRO | FIN-RYT | 0.767718 | 0.00363 | 211.488 | Direct estimation of EL ancestries |
| FIN-PUL | PUN-TYM | SWE-BOL | GBR-GRO | FIN-RYT | 0.779256 | 0.003502 | 222.498 | Direct estimation of EL ancestries |
| FIN-PUL | PUN-TYM | FIN-KIV | GBR-GRO | FIN-RYT | 0.792096 | 0.003531 | 224.338 | Direct estimation of EL ancestries |
| FIN-PUL | PUN-TYM | SWE-BYN | GBR-GRO | FIN-RYT | 0.934234 | 0.004298 | 217.344 | Direct estimation of EL ancestries |
| FIN-PUL | PUN-TYM | SWE-KIR | GBR-GRO | FIN-RYT | 0.983444 | 0.003706 | 265.372 | Direct estimation of EL ancestries |
| FIN-PUL | PUN-TYM | FIN-KAR | GBR-GRO | FIN-RYT | 1.059394 | 0.003642 | 290.872 | Direct estimation of EL ancestries |

| | | | | | | | | |
|---|---|---|---|---|---|---|---|---|
| RUS-BOL | PUN-TYM | DEN-NOR | GBR-GRO | FIN-RYT | 0.250395 | 0.003664 | 68.342 | Direct estimation of EL ancestries |
| RUS-BOL | PUN-TYM | SWE-FIS | GBR-GRO | FIN-RYT | 0.436784 | 0.004192 | 104.206 | Direct estimation of EL ancestries |
| RUS-BOL | PUN-TYM | BEL-MAL | GBR-GRO | FIN-RYT | 0.191226 | 0.003875 | 49.349 | Direct estimation of EL ancestries |
| RUS-BOL | PUN-TYM | GER-RUE | GBR-GRO | FIN-RYT | 0.612494 | 0.00362 | 169.21 | Direct estimation of EL ancestries |
| RUS-BOL | PUN-TYM | FIN-HEL | GBR-GRO | FIN-RYT | 0.784351 | 0.002915 | 269.064 | Direct estimation of EL ancestries |
| RUS-BOL | PUN-TYM | SWE-BOL | GBR-GRO | FIN-RYT | 0.791818 | 0.002788 | 283.974 | Direct estimation of EL ancestries |
| RUS-BOL | PUN-TYM | FIN-KIV | GBR-GRO | FIN-RYT | 0.799974 | 0.002865 | 279.178 | Direct estimation of EL ancestries |
| RUS-BOL | PUN-TYM | SWE-BYN | GBR-GRO | FIN-RYT | 0.932916 | 0.003198 | 291.71 | Direct estimation of EL ancestries |
| RUS-BOL | PUN-TYM | SWE-KIR | GBR-GRO | FIN-RYT | 0.966409 | 0.002745 | 352.048 | Direct estimation of EL ancestries |
| RUS-BOL | PUN-TYM | FIN-KAR | GBR-GRO | FIN-RYT | 0.991859 | 0.002211 | 448.699 | Direct estimation of EL ancestries |
| RUS-BOL | PUN-TYM | DEN-NOR | GBR-GRO | FIN-PUL | 0.249741 | 0.003616 | 69.062 | Direct estimation of EL ancestries |
| RUS-BOL | PUN-TYM | SWE-FIS | GBR-GRO | FIN-PUL | 0.435671 | 0.004131 | 105.465 | Direct estimation of EL ancestries |
| RUS-BOL | PUN-TYM | BEL-MAL | GBR-GRO | FIN-PUL | 0.190725 | 0.003853 | 49.504 | Direct estimation of EL ancestries |
| RUS-BOL | PUN-TYM | GER-RUE | GBR-GRO | FIN-PUL | 0.610936 | 0.003526 | 173.271 | Direct estimation of EL ancestries |
| RUS-BOL | PUN-TYM | FIN-HEL | GBR-GRO | FIN-PUL | 0.782369 | 0.002792 | 280.221 | Direct estimation of EL ancestries |
| RUS-BOL | PUN-TYM | SWE-BOL | GBR-GRO | FIN-PUL | 0.789821 | 0.002656 | 297.323 | Direct estimation of EL ancestries |
| RUS-BOL | PUN-TYM | FIN-KIV | GBR-GRO | FIN-PUL | 0.797961 | 0.002664 | 299.548 | Direct estimation of EL ancestries |
| RUS-BOL | PUN-TYM | SWE-BYN | GBR-GRO | FIN-PUL | 0.930651 | 0.003135 | 296.815 | Direct estimation of EL ancestries |
| RUS-BOL | PUN-TYM | SWE-KIR | GBR-GRO | FIN-PUL | 0.964043 | 0.002392 | 402.958 | Direct estimation of EL ancestries |
| RUS-BOL | PUN-TYM | FIN-KAR | GBR-GRO | FIN-PUL | 0.989394 | 0.001821 | 543.291 | Direct estimation of EL ancestries |
| FIN-RYT | PUN-TYM | DEN-NOR | GBR-GRO | FIN-PUL | 0.248912 | 0.003735 | 66.652 | Direct estimation of EL ancestries |
| FIN-RYT | PUN-TYM | SWE-FIS | GBR-GRO | FIN-PUL | 0.431463 | 0.00431 | 100.119 | Direct estimation of EL ancestries |
| FIN-RYT | PUN-TYM | BEL-MAL | GBR-GRO | FIN-PUL | 0.190848 | 0.003924 | 48.63 | Direct estimation of EL ancestries |
| FIN-RYT | PUN-TYM | GER-RUE | GBR-GRO | FIN-PUL | 0.608269 | 0.003885 | 156.554 | Direct estimation of EL ancestries |
| FIN-RYT | PUN-TYM | FIN-HEL | GBR-GRO | FIN-PUL | 0.783608 | 0.00348 | 225.166 | Direct estimation of EL ancestries |
| FIN-RYT | PUN-TYM | SWE-BOL | GBR-GRO | FIN-PUL | 0.790574 | 0.003347 | 236.218 | Direct estimation of EL ancestries |
| FIN-RYT | PUN-TYM | FIN-KIV | GBR-GRO | FIN-PUL | 0.797618 | 0.003356 | 237.664 | Direct estimation of EL ancestries |
| FIN-RYT | PUN-TYM | SWE-BYN | GBR-GRO | FIN-PUL | 0.953706 | 0.004619 | 206.458 | Direct estimation of EL ancestries |
| FIN-RYT | PUN-TYM | SWE-KIR | GBR-GRO | FIN-PUL | 0.981609 | 0.003547 | 276.723 | Direct estimation of EL ancestries |
| FIN-RYT | PUN-TYM | FIN-KAR | GBR-GRO | FIN-PUL | 0.999289 | 0.002769 | 360.885 | Direct estimation of EL ancestries |
| GBR-GRO | PUN-TYM | DEN-NOR | RUS-BOL | BEL-MAL | 0.880991 | 0.002722 | 323.626 | Direct estimation of WL ancestries |
| GBR-GRO | PUN-TYM | SWE-FIS | RUS-BOL | BEL-MAL | 0.547724 | 0.005187 | 105.593 | Direct estimation of WL ancestries |

| | | | | | | | | |
|---|---|---|---|---|---|---|---|---|
| GBR-GRO | PUN-TYM | GER-RUE | RUS-BOL | BEL-MAL | 0.301761 | 0.004264 | 70.763 | Direct estimation of WL ancestries |
| GBR-GRO | PUN-TYM | FIN-HEL | RUS-BOL | BEL-MAL | 0.113691 | 0.003287 | 34.586 | Direct estimation of WL ancestries |
| GBR-GRO | PUN-TYM | SWE-BOL | RUS-BOL | BEL-MAL | 0.108377 | 0.003052 | 35.508 | Direct estimation of WL ancestries |
| GBR-GRO | PUN-TYM | FIN-KIV | RUS-BOL | BEL-MAL | 0.102107 | 0.003093 | 33.015 | Direct estimation of WL ancestries |
| GBR-GRO | PUN-TYM | SWE-BYN | RUS-BOL | BEL-MAL | 0.026316 | 0.003146 | 8.364 | Direct estimation of WL ancestries |
| GBR-GRO | PUN-TYM | SWE-KIR | RUS-BOL | BEL-MAL | 0.012801 | 0.002706 | 4.731 | Direct estimation of WL ancestries |
| GBR-GRO | PUN-TYM | FIN-KAR | RUS-BOL | BEL-MAL | 0.006021 | 0.002009 | 2.997 | Direct estimation of WL ancestries |
| GBR-GRO | PUN-TYM | DEN-NOR | FIN-PUL | BEL-MAL | 0.880462 | 0.002731 | 322.342 | Direct estimation of WL ancestries |
| GBR-GRO | PUN-TYM | SWE-FIS | FIN-PUL | BEL-MAL | 0.545745 | 0.005336 | 102.272 | Direct estimation of WL ancestries |
| GBR-GRO | PUN-TYM | GER-RUE | FIN-PUL | BEL-MAL | 0.29868 | 0.004421 | 67.567 | Direct estimation of WL ancestries |
| GBR-GRO | PUN-TYM | FIN-HEL | FIN-PUL | BEL-MAL | 0.109778 | 0.00351 | 31.278 | Direct estimation of WL ancestries |
| GBR-GRO | PUN-TYM | SWE-BOL | FIN-PUL | BEL-MAL | 0.104445 | 0.00327 | 31.945 | Direct estimation of WL ancestries |
| GBR-GRO | PUN-TYM | FIN-KIV | FIN-PUL | BEL-MAL | 0.098143 | 0.003205 | 30.624 | Direct estimation of WL ancestries |
| GBR-GRO | PUN-TYM | SWE-BYN | FIN-PUL | BEL-MAL | 0.022032 | 0.003416 | 6.449 | Direct estimation of WL ancestries |
| GBR-GRO | PUN-TYM | SWE-KIR | FIN-PUL | BEL-MAL | 0.008443 | 0.002812 | 3.002 | Direct estimation of WL ancestries |
| GBR-GRO | PUN-TYM | FIN-KAR | FIN-PUL | BEL-MAL | 0.001652 | 0.002324 | 0.711 | Direct estimation of WL ancestries |
| GBR-GRO | PUN-TYM | DEN-NOR | FIN-RYT | BEL-MAL | 0.880531 | 0.002742 | 321.099 | Direct estimation of WL ancestries |
| GBR-GRO | PUN-TYM | SWE-FIS | FIN-RYT | BEL-MAL | 0.545979 | 0.005283 | 103.35 | Direct estimation of WL ancestries |
| GBR-GRO | PUN-TYM | GER-RUE | FIN-RYT | BEL-MAL | 0.299071 | 0.004415 | 67.735 | Direct estimation of WL ancestries |
| GBR-GRO | PUN-TYM | FIN-HEL | FIN-RYT | BEL-MAL | 0.110277 | 0.003527 | 31.268 | Direct estimation of WL ancestries |
| GBR-GRO | PUN-TYM | SWE-BOL | FIN-RYT | BEL-MAL | 0.104939 | 0.003291 | 31.884 | Direct estimation of WL ancestries |
| GBR-GRO | PUN-TYM | FIN-KIV | FIN-RYT | BEL-MAL | 0.09865 | 0.003375 | 29.228 | Direct estimation of WL ancestries |
| GBR-GRO | PUN-TYM | SWE-BYN | FIN-RYT | BEL-MAL | 0.02261 | 0.003396 | 6.658 | Direct estimation of WL ancestries |
| GBR-GRO | PUN-TYM | SWE-KIR | FIN-RYT | BEL-MAL | 0.00898 | 0.003259 | 2.755 | Direct estimation of WL ancestries |
| GBR-GRO | PUN-TYM | FIN-KAR | FIN-RYT | BEL-MAL | 0.002187 | 0.002584 | 0.846 | Direct estimation of WL ancestries |
| NOTE- Orange colored rows marks the data used to generate Fig.2 (c) for the sake of clarity. | | | | | | | | |

Table S5. Result of f4-ratio test and enrichment or depletion of ancestry proportion (alpha) in different genomic categories of admixed populations.

| A | O | X | C | B | alpha | std | z | feature | manner | *p* alpha higher | p alpha lower |
|---|---|---|---|---|---|---|---|---|---|---|---|
| FIN-PUL | PUN-TYM | DEN-NOR | GBR-GRO | RUS-BOL | 0.25662 | 0.00444 | 57.781 | Intergenic | Direct estimation of EL ancestries | | |
| FIN-PUL | PUN-TYM | BEL-MAL | GBR-GRO | RUS-BOL | 0.19987 | 0.00469 | 42.601 | Intergenic | Direct estimation of EL ancestries | | |
| GBR-GRO | PUN-TYM | FIN-HEL | RUS-BOL | BEL-MAL | 0.11298 | 0.00399 | 28.327 | Intergenic | Direct estimation of WL ancestries | | |
| GBR-GRO | PUN-TYM | SWE-BOL | RUS-BOL | BEL-MAL | 0.10831 | 0.004 | 27.076 | Intergenic | Direct estimation of WL ancestries | | |
| GBR-GRO | PUN-TYM | FIN-KIV | RUS-BOL | BEL-MAL | 0.10237 | 0.00403 | 25.405 | Intergenic | Direct estimation of WL ancestries | | |
| FIN-PUL | PUN-TYM | DEN-NOR | GBR-GRO | RUS-BOL | 0.23162 | 0.00632 | 36.661 | CDS | Direct estimation of EL ancestries | 0.9999 | 0.000001 |
| FIN-PUL | PUN-TYM | BEL-MAL | GBR-GRO | RUS-BOL | 0.16701 | 0.00662 | 25.224 | CDS | Direct estimation of EL ancestries | 0.9999 | 0.000001 |
| GBR-GRO | PUN-TYM | FIN-HEL | RUS-BOL | BEL-MAL | 0.12347 | 0.00561 | 22.015 | CDS | Direct estimation of WL ancestries | 0.0274 | 0.972503 |
| GBR-GRO | PUN-TYM | SWE-BOL | RUS-BOL | BEL-MAL | 0.1155 | 0.00537 | 21.53 | CDS | Direct estimation of WL ancestries | 0.08809 | 0.911809 |
| GBR-GRO | PUN-TYM | FIN-KIV | RUS-BOL | BEL-MAL | 0.1073 | 0.00528 | 20.34 | CDS | Direct estimation of WL ancestries | 0.18338 | 0.816518 |
| FIN-PUL | PUN-TYM | DEN-NOR | GBR-GRO | RUS-BOL | 0.25599 | 0.00863 | 29.666 | ConstrainedElements | Direct estimation of EL ancestries | 0.52385 | 0.476052 |
| FIN-PUL | PUN-TYM | BEL-MAL | GBR-GRO | RUS-BOL | 0.20017 | 0.00882 | 22.69 | ConstrainedElements | Direct estimation of EL ancestries | 0.48825 | 0.511649 |
| GBR-GRO | PUN-TYM | FIN-HEL | RUS-BOL | BEL-MAL | 0.10097 | 0.00624 | 16.193 | ConstrainedElements | Direct estimation of WL ancestries | 0.9718 | 0.028097 |
| GBR-GRO | PUN-TYM | SWE-BOL | RUS-BOL | BEL-MAL | 0.09835 | 0.006 | 16.388 | ConstrainedElements | Direct estimation of WL ancestries | 0.9539 | 0.045995 |
| GBR-GRO | PUN-TYM | FIN-KIV | RUS-BOL | BEL-MAL | 0.08716 | 0.00595 | 14.641 | ConstrainedElements | Direct estimation of WL ancestries | 0.9946 | 0.005299 |
| FIN-PUL | PUN-TYM | DEN-NOR | GBR-GRO | RUS-BOL | 0.24721 | 0.00435 | 56.813 | Intron | Direct estimation of EL ancestries | 0.9859 | 0.013999 |
| FIN-PUL | PUN-TYM | BEL-MAL | GBR-GRO | RUS-BOL | 0.18983 | 0.00454 | 41.792 | Intron | Direct estimation of EL ancestries | 0.9863 | 0.013599 |
| GBR-GRO | PUN-TYM | FIN-HEL | RUS-BOL | BEL-MAL | 0.11091 | 0.00379 | 29.288 | Intron | Direct estimation of WL ancestries | 0.71103 | 0.288871 |
| GBR-GRO | PUN-TYM | SWE-BOL | RUS-BOL | BEL-MAL | 0.10548 | 0.00357 | 29.576 | Intron | Direct estimation of WL ancestries | 0.78352 | 0.216378 |
| GBR-GRO | PUN-TYM | FIN-KIV | RUS-BOL | BEL-MAL | 0.09947 | 0.00344 | 28.948 | Intron | Direct estimation of WL ancestries | 0.79922 | 0.20068 |
| FIN-PUL | PUN-TYM | DEN-NOR | GBR-GRO | RUS-BOL | 0.23335 | 0.00718 | 32.498 | Promoter | Direct estimation of EL ancestries | 0.9993 | 0.0006 |
| FIN-PUL | PUN-TYM | BEL-MAL | GBR-GRO | RUS-BOL | 0.18254 | 0.00741 | 24.637 | Promoter | Direct estimation of EL ancestries | 0.9894 | 0.010499 |
| GBR-GRO | PUN-TYM | FIN-HEL | RUS-BOL | BEL-MAL | 0.12643 | 0.00673 | 18.776 | Promoter | Direct estimation of WL ancestries | 0.023 | 0.976902 |
| GBR-GRO | PUN-TYM | SWE-BOL | RUS-BOL | BEL-MAL | 0.11959 | 0.00668 | 17.91 | Promoter | Direct estimation of WL ancestries | 0.0477 | 0.952205 |
| GBR-GRO | PUN-TYM | FIN-KIV | RUS-BOL | BEL-MAL | 0.11434 | 0.00663 | 17.254 | Promoter | Direct estimation of WL ancestries | 0.0377 | 0.962204 |

Table S6. Test of ILS and detailed information for each candidate region identified by fd analysis.

| Chr | Start | End | Length | Structural Variant | Mean recombi | p(0.43) | p(0.67) | p(0.92) | L(0.43) | L(0.67) | L(0.92) | T(exp) | Genes |
|---|---|---|---|---|---|---|---|---|---|---|---|---|---|
| LG1 | 8880001 | 9040000 | 159306 | | 4.28E-10 | 6.71E-06 | 2.85E-09 | 7.67E-13 | 10,864.03 | 6,972.44 | 5,077.75 | 29,324.27 | RUNX1; DONSON; Arhgap1; Arl6ip6; SLC19A2; ATG3; CARS2; DDRGK1 |
| LG1 | 25740001 | 25920000 | 178684 | Dup. | 1.15E-07 | 0 | 0 | 0 | 40.44 | 25.95 | 18.90 | 97.32 | MMP20; MMP13; TSKU; GUCY2F; SERPINH1; GDPD5; HSPA13; SAMSN1; ZP4; P2RX5; Dhx40; AKT2; CNTD2; CIPC; SLC6A13; Vwa7; VWA5A |
| LG1 | 28680001 | 28880000 | 199904 | Dup. | 4.89E-09 | 0 | 0 | 0 | 951.16 | 610.44 | 444.56 | 2,045.97 | NLK; SMCO4; NF1; SWP2; COX7A2; Ribonuclease Oy; SUPT5H; TRIAP1; IL15; Plekhg3; P2RY14; SHKBP1; Ltbp2; Ltbp4 |
| LG3 | 2180001 | 2280000 | 90059 | | 4.30E-08 | 0 | 0 | 0 | 108.14 | 69.40 | 50.54 | 516.34 | RNF31; INHBA; Amph; Acot5; ACOT1 |
| LG3 | 15520001 | 15640000 | 119478 | | 3.17E-09 | 0 | 0 | 0 | 1,468.79 | 942.66 | 686.50 | 5,286.15 | Lyn; PKIA; Ankrd33b; Xrn1; TOE1; Prkrir; Sepp1b; ELOVL4; FABP6 |
| LG4 | 23360001 | 23460000 | 99928 | | 1.00E-10 | 0.367 | 0.153 | 0.056 | 46,511.63 | 29,850.75 | 21,739.13 | 200,144.10 | Pol; Arsi; Tmco6; Aldob; RNF20 |
| LG4 | 24400001 | 24500000 | 91406 | | 1.00E-10 | 0.415 | 0.19 | 0.077 | 46,511.63 | 29,850.75 | 21,739.13 | 218,804.02 | TRPC7 |
| LG8 | 4840001 | 4980000 | 139756 | Dup. | 2.09E-08 | 0 | 0 | 0 | 222.96 | 143.10 | 104.21 | 686.01 | GPX7; SPBPJ4664.02; Arhgap42a |
| LG8 | 13640001 | 13740000 | 99814 | Dup. & Del. | 3.36E-08 | 0 | 0 | 0 | 138.51 | 88.89 | 64.74 | 596.70 | P3H2; Ccdc181; CPOX; PRRG1; TMEM47 |
| LG9 | 2680001 | 2840000 | 159796 | Del. | 6.80E-08 | 0 | 0 | 0 | 68.42 | 43.91 | 31.98 | 184.12 | RASD2; PRPSAP1; PIK3R5; PIK3R6; NTN1; STX8 |
| LG9 | 11760001 | 11860000 | 98740 | Dup. | 6.48E-09 | 0 | 0 | 0 | 717.77 | 460.66 | 335.48 | 3,125.80 | WDR87; ADRA1D; Taar1; AHNAK; Sf3a2 |
| LG10 | 4320001 | 4420000 | 99184 | Dup. & Del. | 8.28E-08 | 0 | 0 | 0 | 56.21 | 36.07 | 26.27 | 243.68 | Slc6a19; Fhod3; Glutathione S-transferase A; KLKB1; F11 |
| LG11 | 8000001 | 8160000 | 159829 | Del. | 3.22E-08 | 0 | 0 | 0 | 144.24 | 92.57 | 67.41 | 388.05 | SSTR2; Usp22; Med9; RASD1; Sox8 |
| LG11 | 14580001 | 14680000 | 99932 | | 2.13E-08 | 0 | 0 | 0 | 218.36 | 140.14 | 102.06 | 939.61 | CLN3; MCT7; DUS1L; SLC16A3 |
| LG11 | 15360001 | 15560000 | 192731 | | 3.97E-08 | 0 | 0 | 0 | 117.08 | 75.14 | 54.72 | 261.22 | |
| LG11 | 17720001 | 17860000 | 120139 | Dup. | 5.19E-08 | 0 | 0 | 0 | 89.60 | 57.51 | 41.88 | 320.71 | SOX10; POLR2F; MICALL1; GGA1; Ddx17; Tnrc6b; Card10; CDC42EP1; Gvin1; GVINP1; F5 |
| LG13 | 17180001 | 17340000 | 159858 | Dup. * | 6.58E-08 | 0 | 0 | 0 | 70.70 | 45.37 | 33.04 | 190.17 | Gastrula zinc finger protein XlCGF57.1; ZNF879; CCDC84 |
| LG13 | 20060001 | 20160000 | 98625 | | 4.27E-08 | 0 | 0 | 0 | 108.96 | 69.93 | 50.93 | 475.08 | TTN; UNC13B |
| LG14 | 2420001 | 2560000 | 139776 | Del. | 8.18E-09 | 0 | 0 | 0 | 568.95 | 365.15 | 265.92 | 1,750.29 | CGNL1; NXPE3; ZP3; CNTFR; Cst4; RASSF6 |
| LG16 | 3020001 | 3120000 | 99660 | | 1.77E-08 | 0 | 0 | 0 | 263.45 | 169.08 | 123.13 | 1,136.69 | Taar1; Taar4 |
| LG18 | 2960001 | 3140000 | 179786 | Inv. ** | 1.26E-07 | 0 | 0 | 0 | 36.86 | 23.65 | 17.23 | 88.15 | Rps29; MGAT2; WDR20; HSP90AB3P; Ppp2r5c; Dio3; Slc25a47a; SLC25A29; Vrtn; Syndig1l |
| LG19 | 20260001 | 20400000 | 105209 | | 4.58E-09 | 0 | 0 | 0 | 1,016.14 | 652.15 | 474.94 | 4,153.08 | Wwox; Maf; F5 |
| LG20 | 620001 | 720000 | 99906 | | 4.54E-08 | 0 | 0 | 0 | 102.47 | 65.77 | 47.89 | 441.04 | ADORA1; Mcam; APOC2; Apoeb; APOA4; MEP1B; FP1; Mep1b; Tomm40; Ddx6 |
| LG20 | 11020001 | 11140000 | 119335 | | 1.00E-10 | 0.274 | 0.092 | 0.027 | 46,511.63 | 29,850.75 | 21,739.13 | 167,595.42 | ZNF226; Trypsin-3; Mypop; ZNF665; ZFP62; ZNF576; ZNF268; ZIM3; ZNF79; DCBLD1; GMEB1 |
| LG20 | 12540001 | 12720000 | 177878 | | 1.00E-10 | 0.105 | 0.018 | 0.003 | 46,511.63 | 29,850.75 | 21,739.13 | 112,436.61 | Grik2 |

NOTE-Dup. Del. and Inv. refers to introgressed duplicate, deletion and inversion, respectively. p(0.43), p(0.67), p(0.92) refers to probability of distinguishing from ILS assuming a divergence time of 0.43, 0.67 and 0.92 Mya, respectively.

L(0.43), L(0.67), L(0.92) refers to expected length resulting from ILS assuming a divergence time of 0.43, 0.67 and 0.92 Mya, respectively. T(exp) refers to Putative age of a given introgressed block T=1/(LR), assuming observed length equals to L.

* Longest introgressed duplicated region amongst all candidate regions

** False positive inference, due to a potential inversion

Table S7. Candidate regions identified by U and Q95 test, and genes in each region.

| Linkage Group (LG | Start | End | Overlap with candidate region from fd ana | Genes in region |
|---|---|---|---|---|
| LG1 | 25740001 | 25860000 | Yes | MMP20; MMP13; TSKU; GUCY2F; SERPINH1; GDPD5; HSPA13; SAMSN1; ZP4; P2RX5; Dhx40;AKT2 |
| LG1 | 28680001 | 28840000 | Yes | NLK; SMCO4; NF1; SWP2; COX7A2; Ribonuclease Oy; SUPT5H; TRIAP1; IL15; PLEKHG3; P2RY14 |
| LG3 | 4940001 | 5120000 | No | Glucagon family neuropeptides; YES1; CLUL1; COLEC12; STAM; TMEM236; CACNB2; MRC1; MALRD1; FKBP1A; PLXDC2; MYRIP |
| LG9 | 2660001 | 2840000 | Yes | RASD2; PRPSAP1; PIK3R5, PIK3R6; NTN1; STX8 |
| LG9 | 4820001 | 5000000 | No | PCDH7; CRMP1; WFS1; PPP2R2C; PDE5A; zgc:101819; ANK2; CAMK2D2 |
| LG11 | 7980001 | 8140000 | Yes | SSTR2; USP22; MED9; RASD1 |
| LG19 | 18120001 | 18220000 | No | MRPL23; CPT1A; SITS-binding protein |

Table S8. Classification of SNPs in NBS (FIN-HEL, SWE-BOL, FIN-KIV) and SWE-BYN population.

| **Focal population** | **Total polymorphic sites** | **Shared with the two source population** | **WL-focal shared *** | | **EL-focal shared *** | | **Unique in focal population** | **Fix between WL and EL** |
|---|---|---|---|---|---|---|---|---|
| | | | ***n*** | ***n* Positively selected** | ***n*** | ***n* Positively selected** | | |
| NBS | 2,776,743 | 636,267 | 331,078 | 518 | 452,613 | 4,787 | 1,356,785 | 83,057 |
| SWE-BYN | 621,771 | 345,432 | 29,102 | | 160,890 | | 86,347 | |

* WL and EL refers to GBR-GRO and RUS-BOL population, separately.

Table S9. Spearman rank correlations (rs) between admixture proportion (fd) and recombination rate for each admixed population.

| **Population** | **Spearman's Correlation test** | |
|---|---|---|
| | ***rs*** | ***p*** |
| SWE-FIS | -0.03 | 0.09 |
| GER-RUE | -0.01 | 0.57 |
| BEL-MAL | -0.17 | <0.001 |
| DEN-NOR | -0.1 | <0.001 |
| FIN-HEL | -0.11 | <0.001 |
| SWE-BOL | -0.13 | <0.001 |
| FIN-KIV | -0.13 | <0.001 |
| SWE-BYN | -0.14 | <0.001 |
| FIN-KAR | -0.19 | <0.001 |
| SWE-KIR | -0.2 | <0.001 |

Table S10. Placement of the randomly selected representative individuals within the phylogenetic trees.

| Individual | Unclassified trees | EL branch-off | WL branch-off |
|---|---|---|---|
| SWE-FIS-35 | 146 | 382 | 469 |
| DEN-NOR-10 | 146 | 134 | 717 |
| BEL-MAL-F26 | 152 | 106 | 739 |
| GBR-GRO-1 | 230 | 28 | 739 |
| GER-RUE-3 | 145 | 605 | 247 |
| 34-m-1 (FIN-HEL) | 140 | 730 | 127 |
| SWE-BOL-69 | 141 | 743 | 113 |
| FIN-KIV-1 | 145 | 750 | 102 |
| FIN-KAR-15 | 135 | 856 | 6 |
| SWE-BYN-1 | 137 | 830 | 30 |
| SWE-KIR-8 | 129 | 851 | 17 |
| FIN-RYT-1 | 110 | 871 | 16 |
| RUS-BOL-33 | 107 | 871 | 19 |
| FIN-PUL-1 | 110 | 871 | 16 |
